# Life, the universe, and everything for $42: ultra-low pass sequencing of maize for genotyping, mapping, and pedigree analysis

**DOI:** 10.64898/2026.02.20.707111

**Authors:** Rajdeep S. Khangura, Amanpreet Kaur, Phillip San Miguel, Brian P. Dilkes

**Affiliations:** Department of Plant and Agroecosystem Sciences, University of Wisconsin-Madison, Madison, WI 53706; Department of Biochemistry, Purdue University, West Lafayette, IN 47097; Center for Plant Biology, Purdue University, West Lafayette, IN 47097; Genomics Facility, Purdue University, West Lafayette, IN 47097

**Keywords:** Mendelian traits, next-generation sequencing, skim sequencing, haplotype detection, identity-by-state, haplotype map, trait integration, background selection, foreground selection

## Abstract

Convenient and economical genotyping methods and simplified bioinformatic workflows are critical for genetic studies and breeding. The declining cost of sequencing library construction, sample multiplexing, and the advent of skim sequencing has reduced costs and enabled large-scale genetic and genomic experiments. Here, we present a simple skim sequencing and bioinformatics pipeline sufficient for various genotyping applications. Our low-depth skim sequencing method costs 21 USD per sample and provides an average of 144k reads. Our approach uses a double-stranded DNA sample to prepare libraries for genome sequencing. We demonstrate various uses for this strategy in maize, a complex and large (2.5 Gbp) genome. DNA from multiple pedigreed populations, including advanced backcrossed progenies, bi-parental populations, near-isogenic lines, and recombinant inbred lines, were used to map loci, detect donor introgressions, and determine introgression haplotypes. Read counts at known polymorphic positions detected donor genotypes even when derived from parents of unknown origin and could localize mutations of phenotypic impact via bulked segregant analysis. Remarkably, the small amount of sequencing data produced were sufficient to identify the haplotypes of introgressions of unknown origin from by comparison to known genotypes. Correct haplotype identification enabled more accurate allele frequencies to be calculated when mapping loci. This is of exceptional value in maize, where a rich collection of mutants from the 20^th^ century are of unknown pedigree.

**One sentence summary:** Economical and efficient whole genome ultra-low pass sequencing of DNA samples for numerous genetic and genomic applications.

## Introduction

The advancement in DNA sequencing technology has resulted in a price reduction in short-read sequencing that resembles the technological advancement in integrated circuits predicted by Moore’s law (Moore 1965). The price of sequencing a human genome has dropped from $10 million USD in 2001 to well below $1000 USD by 2022 (Wetterstrand 2024). This has permitted sequencing large populations of humans, fruit flies, maize, and other species (Auton et al. 2015; Lack et al. 2016; Bukowski et al. 2018). As a result of the shift in costs, experimental designs that apply sequencing to pre-existing pedigrees and reconsider the relative efforts of sequencing and population construction are timely.

Further sequencing cost reductions come from experimental strategies that target specific genomic regions. Among these approaches, genome reduction (Baird et al. 2008; Elshire et al. 2011; Stolle and Moritz 2013) and target-enrichment (Turner et al. 2009; Mamanova et al. 2010) are the widely used for population-level genotyping. These genotyping-by-sequencing (GBS) approaches produce sequence data from a subset of genomic regions across samples for genetic analysis. For instance, the DNA flanking a five-base-pair restriction endonuclease recognition sequence was sequenced in maize and yielded ∼800k sequence tags (Elshire et al. 2011). Adding a second restriction endonuclease to make a more selective GBS strategy, using six and four-base pair recognition sequences, resulted in 200k-400k sequence tags in the larger Barley and Wheat genomes (Poland et al. 2012). Reduced representation sequencing requires special sample preparation steps and some target enrichment approaches require reagents specific to a particular experiment or organism (Jupe et al. 2013; Shih et al. 2018). As sequencing costs decrease, cost savings may not occur or may not be worth the added complexity. Adoption of simple, inexpensive, low-data-per-sample approaches could accelerate genetic research and enable trainees and smaller commercial enterprises to perform advanced genetic and genomic experiments without large initial capital investments.

The Purdue Genomics Facility developed a low-coverage sequencing technique for double-stranded DNA samples that uses the Illumina MiSeq, Element AVITI, or similar mid-scale sequencing approaches. At the time the data in the manuscript was collected, this provided an average of 144,000 reads per sample. This method, called WideSeq, is suitable for high coverage sequencing of plasmids and pooled PCR amplicons and permits *de novo* assembly of products up to ∼100kb. Sequencing PCR amplicons using this approach has previously accomplished a variety of genetic analyses. WideSeq profiling of *Mutator* flanking sequences identified all *Mutator* insertions in a maize mutant population (Zhang et al. 2020). WideSeq of RT-PCR products from a single gene coupled with the analysis of single-nucleotide polymorphisms (SNPs) assessed allele-specific gene expression (Khangura et al. 2019; Zhang et al. 2020). We have also used WideSeq to sequence PCR amplicons from a single locus in maize to determine the molecular nature of somatic mutations caused by ethyl methanesulfonate in chimeric leaves (Karre et al. 2021). We have also used WideSeq to determine the molecular nature of new mutants uncovered by allelism tests (Sauer et al. 2023; Kaur, Best, et al. 2024) and of mutants that mapped to the same location but cannot be tested by allelism due to haploidy (Burow et al. 2024). We have also used low pass whole genome sequencing of EMS mutants to confirm line genotypes and estimate error rates in large scale pedigrees (Addo-Quaye et al. 2018).

Sequencing data is often used in genetic analysis for many goals that do not require data coverage of the entire genome. For example, researchers may (1) map a locus, (2) identify desirable recombinants, (3) genotype germplasm for trait integration, or (4) identify the genetic background of an unknown donor. We implemented an approach that achieves all of these goals using low-coverage sequencing of random genomic regions, provided that validated polymorphisms were already known. This was done with extremely low sequence coverage in maize, which has a large, complex genome, high-quality reference genomes, and robust population genetic resources.

In this study, we demonstrate that 0.01X coverage of whole-genome DNA from maize samples was effective for a variety of downstream genetic and genomic analyses. The sequencing protocol produced an average of 141,000 mapped read pairs (2 x 250 bp) per sample. This low-coverage sequence data was sufficient to perform introgression mapping of maize mutants, detect recombinants, genotype recombinant inbred lines (RILs) and near-isogenic lines (NILs), determine the parent or haplotype-of-origin of classical maize mutants, and identify contaminants in a mutant suppressor screen. This was made possible by the high-quality reference genome and population-wide SNP genotypes from a haplotype map (Bukowski et al. 2018). While donor background was not required, we demonstrated that when donor background is known, low-pass sequencing can achieve greater precision. Since the input was double stranded DNA product, many applications can be envisioned to test a variety of molecular hypotheses. The low data volumes and low compute requirements make this amenable for teaching bioinformatics, as all operations can be executed on a personal computer or laptop.

## Methods

### Plant materials and growth

The *br2-ref* mutant allele described in this work was obtained from the Maize Genetics COOP stock center in a mixed genetic background (stock:114F *br2;hm1;Hm2*). The *br1-ref* allele was obtained from the Maize Genetics COOP stock center in a background of mixed parentage. The b94 and m97 are part of the near-isogenic lines described previously (Eichten et al. 2011), and were obtained from the Maize Genetics COOP stock center (stocks: MBNIL_b094, and MBNIL_m097). The *Sdw2-N1991* allele was originally designated as D*-N1991 and was also obtained from the Maize Genetics COOP stock center (stock# 307A). The *Sdw2-N1991* was obtained in the M1 generation following pollen mutagenesis with nitrosoguanidine (Neuffer 1990; 1992). The NAM-RILs were previously described (Buckler et al. 2009) and were bulk-propagated at Purdue University for use by multiple maize genetics laboratories. We obtained the NAM-RIL accession Z004E0189 from Dr. Mitch Tuinstra (Purdue University).

Plants were either grown in the field at the Purdue Agronomy Center for Research and Education (ACRE) in West Lafayette or in the Purdue Horticulture greenhouse. In the field, each entry was planted in a single-row plot 15 feet long, with 12.5 feet planted and 2.5 feet of walking alley, and row-to-row spacing was fixed at 2.5 feet. The planting density was maintained at ∼15 plants per plot. The standard cultivation practices for growing maize in Indiana were implemented by ACRE staff, including pre-emergence herbicide and fertilizer applications. No supplemental irrigation was needed as adequate rainfall was received throughout the growing season. In the greenhouse, plants were grown under a 16h:8h day:light cycle using high-pressure sodium light as supplemental lighting. Plants were grown in growing media consisting of a ∼1:1 ratio of Berger BM7 bark mix and calcined clay (Turface). The daytime temperature of ∼80° F and nighttime temperature of ∼70° F were maintained in the greenhouse during the duration of plant growth.

### WideSeq library preparation and sequencing

DNA was extracted from leaf tissues using the standard CTAB protocol (Saghai-Maroof et al. 1984). DNA concentrations were quantified using NanoDrop 2000. The method uses a double-stranded DNA sample (e.g., whole-genomic DNA, plasmids, or PCR products) for library preparation with the Tn5-based Nextera Flex DNA Library Preparation Kit from Illumina. The libraries containing adapter and barcode sequences were loaded onto the flow cell lane and sequenced on the MiSeq at the Purdue Genomics Facility. After the quality check, the 250 bp paired-end reads were used for downstream bioinformatic analysis. The cost of sequencing one sample was 21 USD for the experiments presented here.

### Sequence alignment, variant detection, and bin frequency calculations

All genomic positions mentioned in this study are B73 version 4, unless specified otherwise. For every known gene discussed in this work, gene models from different assemblies and annotation pipelines of the B73 genome are reported. The bioinformatic analysis performed using the paired-end sequencing reads is as follows. The adapter sequence and low-quality base pairs from paired-end reads were removed using *trimmomatic* with the default settings (Bolger et al. 2014). The high-quality paired-end reads were aligned to the maize B73 v4 genome assembly using *bwa* (Li and Durbin 2009). The deduplicated BAM files were used to call all positions using the *mpileup* function in *bcftools* (Li 2011; Danecek et al. 2021). For HapMap3 (HapMap3) based mapping or genotyping utility, we used *vcftools* (Danecek et al. 2011) to identify positions with reads that overlapped any of the 55.2 million HapMap3 SNPs (Bukowski et al. 2018). For a known background-based genotyping, inbred-specific polymorphic SNPs relative to B73 were identified from HapMap3 SNPs (Bukowski et al. 2018) using *vcftools*.

For allele frequency calculation at the bin level, we first created a bed file with a given bin size (1 Mbp or 5 Mbp) using *bedtools* (Quinlan and Hall 2010). The counts of reference and non-reference reads at each position overlapping the known HapMap3 SNPs or an inbred-specific polymorphic SNPs were tabulated into 1 or 5Mbp bins using the “*map*” function in *bedtools*. The total counts of reads in each bin were used to calculate the reference or non-reference allele frequencies within and between samples.

### Haplotype detection

We first identified the genomic regions of interest in each sample. The non-reference HapMap3 SNP positions in each sample were retained using a custom *awk* command. All HapMap3 positions identified in the first step were used to keep only the genomic region of interest. To minimize the rate of false positive calls in our analysis, the HapMap3 SNPs across all 1210 accessions were filtered to remove positions with minor allele frequency below 0.05. The final Hmp3 SNPs for the heterozygous samples contained only the non-reference SNPs, whereas the homozygous samples contained both reference and non-reference SNPs in the genomic region of interest. The non-reference alleles in the final VCF file were encoded as “1” and reference alleles as “0”. The SNP genotypes in the VCF files were filtered using *vcftools* (Danecek et al. 2011). We calculated the Jaccard Index, also known as the Jaccard similarity coefficient, using pairwise comparisons of each accession in the HapMap3 panel with the test sample (Gilbert 1884; Jaccard 1901). The accession with the highest genetic similarity will have a Jaccard Index close to 1, and the one with the lowest genetic similarity will be near 0.

### Comparison of recombination breakpoints in NILs and RILs

To compare the recombination breakpoints between our sequencing data and previously published genotypes of near-isogenic lines and the NAM RILs, we employed the following approach. The genotypes of the NILs (B73 AGPv2) were obtained from a previous study that described this population (Eichten et al. 2011). The NAM RILs genotypes were obtained from the Panzea project through CyVerse. The NAM-RILs were previously genotyped using the Illumina Golden Gate Assay as described by McMullen *et al*. 2009. The chain file for v2->v4 conversion was obtained from the Gramene database. The B73 RefGenv2 positions were matched to B73 RefGenv4 using CrossMap v0.6.3. The parameter “--no-comp-alleles” was included in CrossMap to place SNP without comparison of the reference allele and the alternate allele. The final count of SNP genotypes imported to B73 AGPv4 coordinates was as follows: For NILs, out of 7204 SNPs, 385 SNPs failed liftover to B73 AGPv4. For NAM RILs, two out of 1028 SNPs failed liftover to B73 AGPv4. The B73 AGPv4 coordinates between the SNP genotypes were compared with the genomic coordinates of the binned allele frequencies calculated from the sequence data approach.

### Data availability

The raw paired-end sequencing data were submitted to NCBI BioProject accession# PRJNA1221219. All the other data, including scripts, are provided with this manuscript.

## Results

### Introgression detection and mapping of recessive maize mutants of unknown parentage by low-pass sequencing

Maize mutants are often generated by EMS mutagenesis in hybrid genetic backgrounds (Candela and Hake 2008; Neuffer et al. 2009). Prior to detailed phenotypic analysis of mutants, the standard practice is to introgress maize mutants into defined genetic backgrounds. Recurrent backcrossing helps to identify backgrounds that provide good mutant expression, remove background mutations, and fix segregating modifiers. As a result, many maize mutants at the stock centers are derived from unknown or mixed genetic backgrounds but available as either advanced backcross generations or mixed parentage multi-generation hybrids. We investigated whether introgressions containing the donor background and causative mutation were distinguishable from the recurrent background with very little sequence data. We attempted to identify introgressed regions in mutants with $21 of sequencing data per sample. Briefly, this approach uses inexpensive *Tn5*-transposon-based “tagmentation” library assembly from double-stranded DNA, followed by sequencing on an Illumina MiSeq instrument (**Figure 1**). Optimized sample numbers and shared sequencing runs across a broad user base (all users of the Purdue University Genomics Core Facility) resulted in a $21 cost for an average of 144,000 250 bp reads per sample.

**Figure 1.**
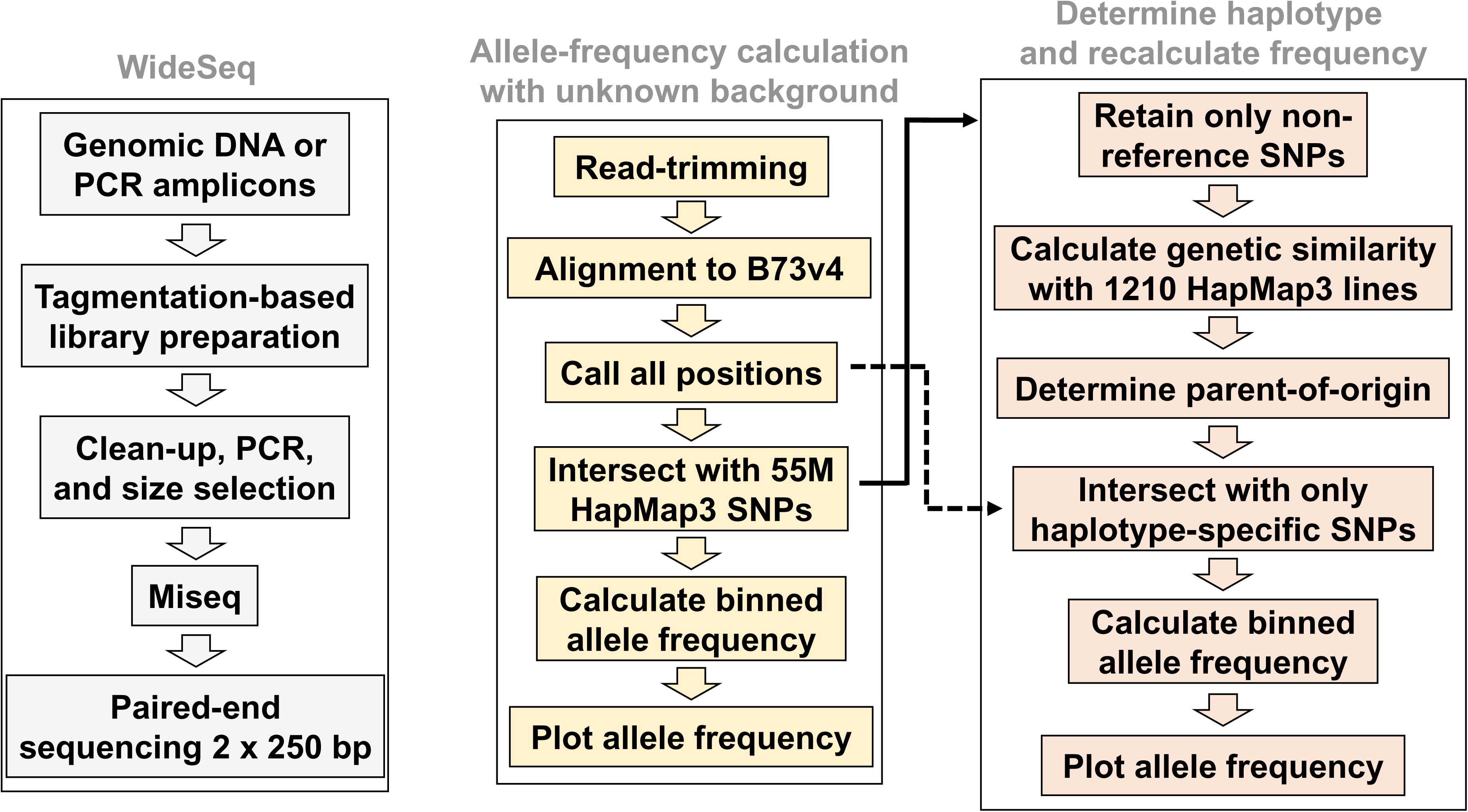
General workflow for (a) WideSeq, (b) allele frequency calculation using WideSeq paired-end reads, and (c) predicting haplotype and recalculating allele frequency using WideSeq paired-end reads

An advanced backcross population of a mutant allele at *brachytic2* (*br2*) into the B73 inbred was used to test the validity of our introgression detection and mapping approach. The molecular identity of *br2* gene is known (Multani et al. 2003). It is located on chr1 (B73 v3: GRMZM2G315375; B73 v4: Zm00001d031871; B73 v5: Zm00001eb038710) and encodes ATP binding cassette type B1. We obtained a recessive *br2-ref* mutant allele derived from an unknown background and linked in cis to a recessive *hm1* allele (Maize Genetics Coop stock#114F *br2;hm1;Hm2*). The *hm1* gene (B73 v3: GRMZM5G881887; B73 v4: Zm00001d031802; B73 v5: Zm00001eb038160) encodes for HC-toxin reductase and provides resistance to a fungal pathogen *Cochliobolus carbonum race1* (Sindhu et al. 2008). The *hm1* gene is ∼2Mb away from the *br2* gene towards the centromere, with the following gene order: *cent1-hm1-br2*. This mutant allele was backcrossed into the B73 background for seven cycles, and then BC7F1 individuals were self-pollinated, and homozygotes were maintained and propagated by self-pollination (**Figure 2**). Recessive *br2* homozygous individuals (n=36) were sampled and DNA extracted as a pool for sequencing. Alignment of the reads to the B73 genome is expected to recover reads with non-reference genotypes from the progenitor background at SNPs encoded at positions linked to the homozygous mutant alleles at *br2-ref* and *hm1*. Low-pass sequencing yielded 257,108 reads that mapped to the B73 v4 reference genome (**Table 1**). We identified all reads that overlapped any of the 55 million HapMap3 SNP positions to genotype all available polymorphisms. HapMap3 SNPs were tabulated from all mapped reads as either missing, reference B73, or non-reference genotypes. The number of reference and non-reference positions were counted in non-overlapping 5Mb genomic bins tiled across the entire genome. The proportion of non-reference genotypes at all observed HapMap3 positions for each bin is graphed in **Figure 2**. The recurrent background, B73, results in a very low frequency of non-reference alleles across most of the genome whereas introgressed segments are clearly visible from the increase in frequency of non-reference reads across multiple contiguous 5Mb bins. As expected, a contiguous stretch of four bins on chr1 with elevated frequencies of non-reference reads is visible across the known positions of *hm1* and *br2* (**Figure 2**). This introgression starts in the bin starting at 200Mb and concludes in the bin ending at 220Mb, spanning the region between *hm1* (∼202.2Mb) and *br2* (∼204.7Mb). The highest non-reference allele frequency, 0.17, was observed in the bin starting at 205Mb and ending at 210Mb (**Figure 2 and Table S1**). We identified two additional contiguous bins on chr9 (140-150Mb) with elevated non-reference allele frequency, demonstrating the near complete conversion of our genetic stock to B73. In addition, using an arbitrary cut-off of non-reference allele frequency of 0.02, there are four single bin-sized departures from expected reference allele frequencies on chr1 (295-300Mb), chr5 (215-220Mb), chr7 (0-5Mb), and chr9 (10-15Mb) (**Figure 2 and Table S1**) that may represent additional introgressed segments or polymorphism between different B73 sources (Liang and Schnable 2016). It is possible that introgressions narrower than 5Mb and regions with little polymorphism between the donor and B73 may give false negative results in this approach. Given that identified 10 bins out of 426 total bins across the maize genome had non-reference allele frequency above 0.02, we estimate that our backcrossed *br2*;*hm1* genetic stock is 98% B73 (**Table S1**) and should be suitable for RNAseq and other detailed characterizations.

**Figure 2.**
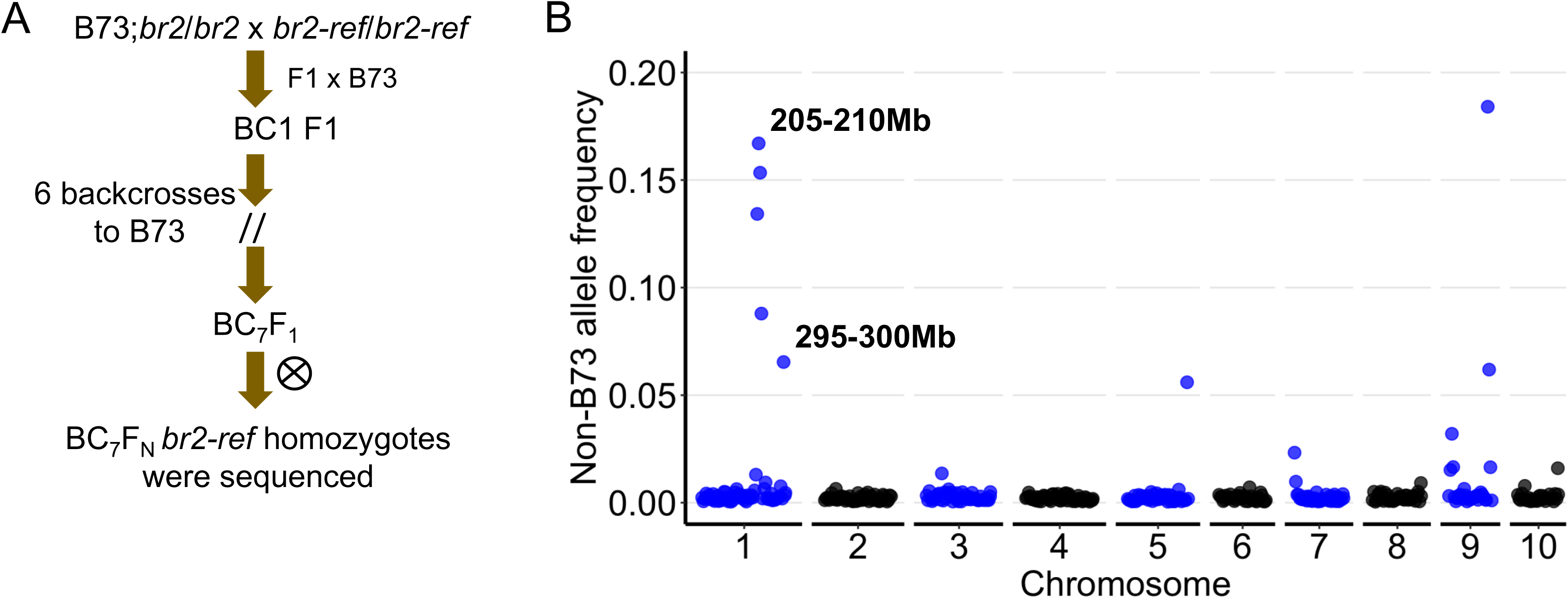
Introgression mapping of *br2-ref* mutant allele. (A) The crossing scheme to derive *br2-ref* introgression into B73 genetic background through recurrent backcrossing. (B) The non-B73 allele frequency in pooled homozygous *br2-ref* individuals sequenced by WideSeq. Each dot represent the 5 Mb genomic intervals with read count at each HapMap3 positions across all ten chromosomes of maize. The start and end position of the top 5Mb bin is annotated.

**Table 1.** Mapping and coverage statistics of all samples described in this study. Average and standard deviation of total QC reads and coverage is provided.

| Sample_ID | QC reads | Mapped Reads | Properly paired reads | Bases mapped | Coverage |
| --- | --- | --- | --- | --- | --- |
| *br2-ref* | 257363 | 257108 | 254156 | 38566200 | 0.016 |
| *br1-ref* | 114227 | 114067 | 113046 | 17110050 | 0.007 |
| b94_NIL | 157808 | 157426 | 154700 | 23613900 | 0.010 |
| *Sdw2-N1991/+* pool | 1284090 | 1269988 | 1243782 | 190498200 | 0.081 |
| *Sdw2-N1991_wt_sib* pool | 1620937 | 1517393 | 1481598 | 227608950 | 0.096 |
| *Sdw2-N1991-sup1* | 160225 | 151458 | 149874 | 22718700 | 0.010 |
| *Sdw2-N1991-sup3* | 97329 | 97077 | 96323 | 14561550 | 0.006 |
| *Sdw2-N1991-sup4* | 105809 | 105548 | 104597 | 15832200 | 0.007 |
| *Sdw2-N1991-sup5* | 132406 | 132133 | 130588 | 19819950 | 0.008 |
| *Sdw2-N1991-sup7* | 103334 | 102907 | 101890 | 15436050 | 0.007 |
| *Sdw2-N1991-sup8* | 225830 | 225456 | 222451 | 33818400 | 0.014 |
| *Sdw2-N1991-sup9* | 176518 | 171519 | 169677 | 25727850 | 0.011 |
| *Sdw2-N1991-sup10* | 127876 | 127709 | 126424 | 19156350 | 0.008 |
| m97_NIL | 83595 | 83254 | 79078 | 12488100 | 0.005 |
| Z004E0189 | 136621 | 136355 | 131694 | 20453250 | 0.009 |
| Average* | 144534 | 143232 | 141115 | 21484812 | 0.009 |
| Standard Deviation | 51096 | 50690 | 50333 | 7603498 | 0.003 |

\**Sdw2-N1991/+* and its wildtype siblings used for Bulk-segregant analysis were excluded from overall summary statistics due to unusually high data output in our WideSeq run.

To identify candidate regions encoding another mutant of known identity, we used a B73 introgression of a mutant allele of *brachytic1* (*br1*) (Kempton 1921). The gene encoding *br1* is linked to *br2* on chr1 (*cent1*-*hm1*-*br2*-*br1*), and encodes a MYB transcription factor (B73 v3: GRMZM2G428555; B73 v4: Zm00001d032194; B73 v5: Zm00001eb041450) (Jiao et al. 2023). We used the original recessive *br1-ref* allele for this experiment which arose spontaneously in traditional populations of maize (Kempton 1921). Using a similar crossing scheme described above for *br2-ref*, we pooled tissue from 36 individuals for low-pass sequencing (**Figure 3)**. The genotypes at polymorphic positions in the full HapMap3 were tabulated for all aligned reads for each 5Mb bin in the same manner as for *br2-ref*. This time $21 of sequencing resulted in 114,067 aligned reads that corresponds to a sequencing depth of 0.007x (**Table 1**). As expected, from the known location of *br1* at ∼216Mb (Jiao et al. 2023), the sequencing data from *br1-ref* homozygotes identified an introgressed segment on chr1 indicated by a high non-reference allele frequency at multiple contiguous bins starting at 200Mb and ending at 220Mb showing peak non-reference allele frequency of 0.12 in the 5Mb bin starting at 215Mb (**Figure** 3). Using an arbitrary threshold of non-reference allele frequency exceeding 0.02, only other bins with non-B73 genomic DNA were identified elsewhere in the *br1-ref* sample, one on chr1 (295-300Mb) and another on chr5 (215-220Mb) (**Figure 3 and Table S2**). Both *br1-ref* and *br2-ref* stocks were backcrossed to the same source of B73 in the lab, and both contain these chr1 and chr5 non-reference bins suggesting that these bins are polymorphic between the B73 used for the introgression (Liang and Schnable 2016) and the reference genome used in HapMap3. The phenotypes of *br1-ref* and *br2-ref* are not distinguishable at the gross morphological level. We confirmed that the material we have are indeed two different mutants through a test cross. As with the *br2-ref* introgression, the *br1-ref* introgression into B73 is nearly complete and ready for additional careful analysis of phenotype. The F1 crosses between *br1-ref* and *br2-ref* complemented their mutant phenotypes confirming that these are two independent but linked genes. Our mapping provides additional molecular evidence that these two mutants, that confer reduced stature phenotype, are linked in maize.

**Figure 3.**
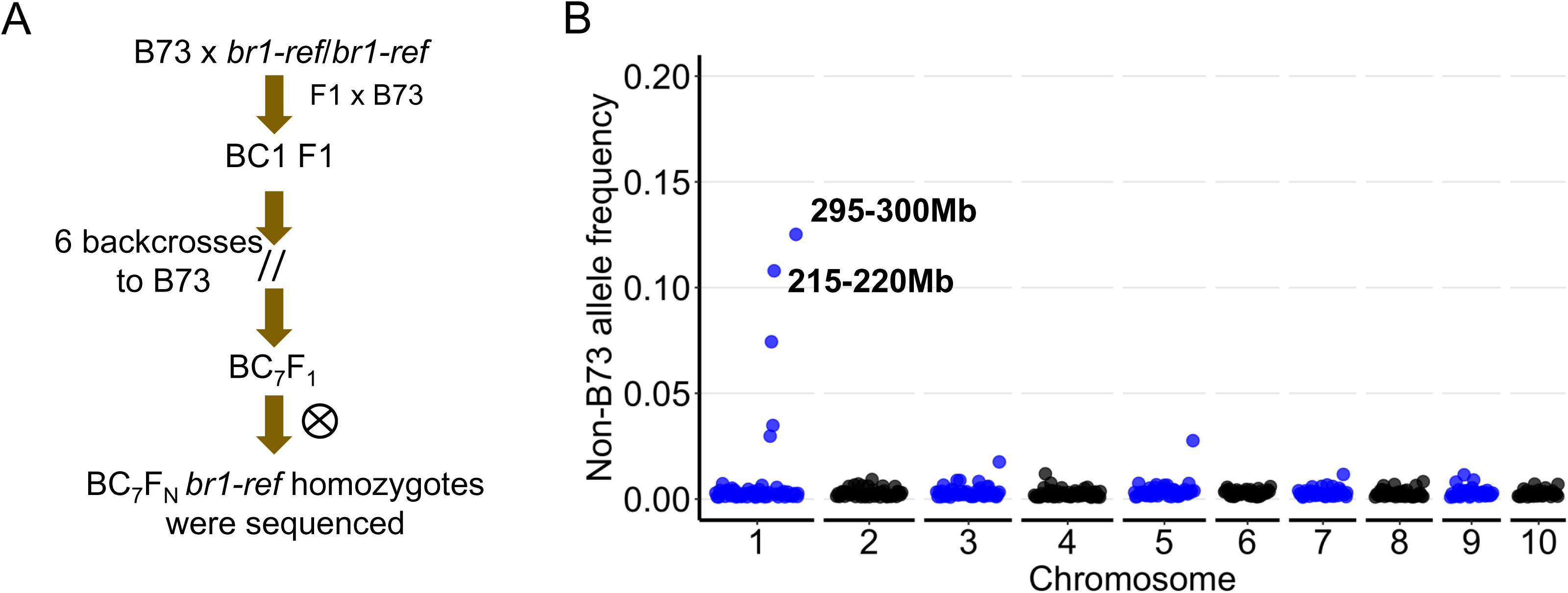
Introgression mapping of *br1-ref* mutant allele. (A) The crossing scheme to derive *br1-ref* introgression into B73 genetic background through recurrent backcrossing. (B) The non-B73 allele frequency in pooled homozygous *br1-ref* individuals sequenced by WideSeq. Each dot represent the 5 Mb genomic intervals with read count at each HapMap3 positions across all ten chromosomes of maize. The start and end position of the top 5Mb bin that contains the *br1* gene is annotated.

### Mapping of a dominant mutant of unknown origin by bulked segregant analysis at low pass for 42 USD

Unlike *br1-ref* and *br2-ref*, many maize mutants are of unknown position. If only a mutant pool is sequenced, any genomic regions with high non-reference allele frequency would represent possible location of the mutant. In those cases, comparison of low-pass sequencing from a pool of mutant individuals to sequence from a pool of wildtype siblings can identify the position of the causative locus (Michelmore et al. 1991). The semi-dominant mutant allele *Sdw2-N1991* is of unknown molecular identity and unknown pedigree that was induced by nitrosoguanidine mutagenesis of pollen (originally called *D*-1991*; Neuffer 1990; Neuffer 1992). This mutant was previously mapped by BA translocation to chr3 and was later determined to be linked to the dominant mutant *Lax midrib1* that is also located on chr3 (Neuffer 1988; 1992; Schichnes et al. 1994; Dotto et al. 2010). A plant height QTL in the NCRPIS Panel was identified at ∼159Mb on chr3 (B73 v4) and *sdw2* was proposed as a candidate for the QTL (Peiffer et al. 2014). To better map the genomic position of the *Sdw2-N1991* allele and test its suitability as a candidate for the plant height QTL, we grew a segregating BC7F1 population from recurrent crosses to B73 (**Figure 4**). We sequenced DNA extracted from pooled leaf tissues from 250 *Sdw2-N1991*/+ plants and 250 wildtype siblings from the BC7F1 population that segregated ∼1:1 for mutant and wildtype siblings (**Figure 4**). The *Sdw2-N1991/+* pool was sequenced at 0.08x depth with a total of 1,269,988 mapped reads, whereas the wildtype pool was sequenced at the 0.09x depth with a total of 1,517,393 mapped reads (**Table 1**). The paired-end reads were aligned to the B73 v4 genome and all sequences overlapping all HapMap3 polymorphic positions were tabulated as reference B73-type or alternative non-B73 type as described above. The non-reference allele frequencies of every 5Mb bin were calculated for *Sdw2-N1991/+* and wildtype sibling pools and graphed (**Figure 4**). This identified a stretch of contiguous bins with a greater proportion of non-B73 allele frequencies in the mutants as compared to wildtype pools on chr3 spanning 15-130Mb (**Figure 4**). The mutant pool had a clear breakpoint where the non-B73 allele frequency increased dramatically in the bin starting at 15 Mb and ended in the bin ending at 125 Mb (**Figure 4 and Table S3**). The wildtype pool, however, showed a clear drop in non-B73 allele frequency from bin starting at 55 Mb and ending at the bin that starts at 110 Mb (**Table S7**). Some genomic regions of non-B73 alleles, as detected in *br1-ref* and *br2-ref,* were also present in both *Sdw2-N1991/+* and wildtype sibling pool on chromosomes 1 (295-300Mb), 4 (20-35Mb), 5 (215-220Mb), and 7 (140-145Mb) (**Figure 4** and **Table S3**). These genomic regions are likely a feature of variation in B73 sources used by different research groups (Liang and Schnable 2016). Based on these data, the gene(s) underlying *Sdw2-N1991* lies within the segment from 15-130 Mb on chr3. This eliminates *sdw2* as a candidate for the maize plant height QTL previously identified at ∼159Mb on chr3 (Peiffer et al. 2014). Thus, bulked segregant analysis at extremely low coverage is an efficient and economical way to map mutants of maize to a defined genomic space that can test candidate genes in linkage mapping studies.

**Figure 4.**
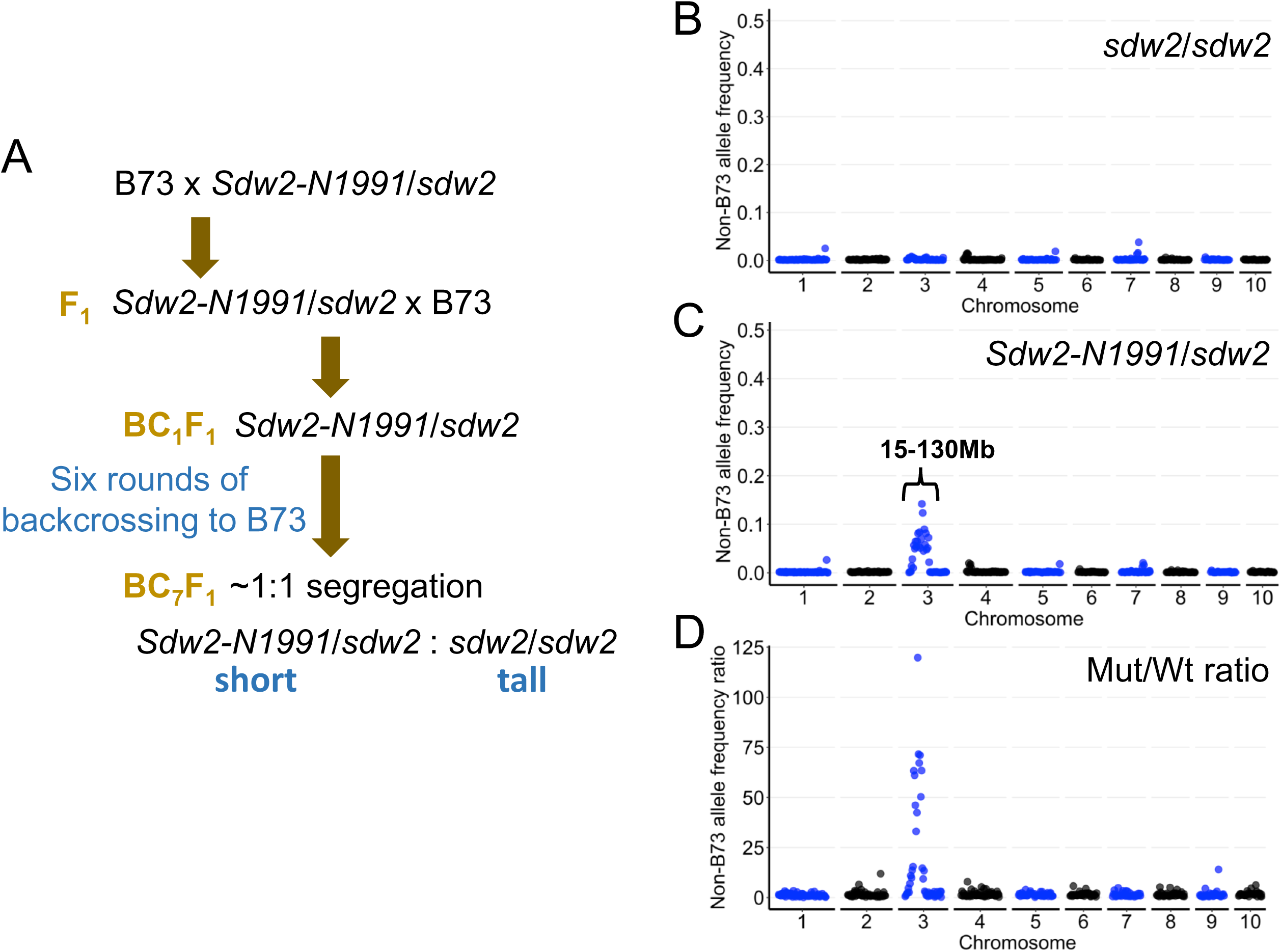
Mapping of *Sdw2-N1991* mutant using bulk-segregant analysis of advanced backcross population. (A) Crossing scheme used to introgress *Sdw2-N1991* allele into B73 background and final samples used for WideSeq. The non-B73 allele frequency at 5Mb bin resolution using WideSeq reads mapped to all hapmap3 positions in the (B) wild-type sibling pool, (C) isogenic *Sdw2-N1991/+* mutant pool. The mutant/wildtype ratio of non-B73 allele frequency in the same 5 Mb bins is plotted in panel D. The start and end position of 5Mb bin with elevated non-B73 allele frequency on chromosome 3 is annotated.

### Using the *Sdw2-N1991* introgression to detect pollen contamination in an EMS-induced suppressor screen by low pass sequencing

The mapping of *Sdw2-N1991* to a defined genomic region on chr3 permitted us to genotype samples by sequencing to rule out any contamination in a suppressor genetic screen. The community standard in maize, and genetic studies in general, is that at least two independent alleles must be identified at a gene to molecularly assign it as the cause of a mutant phenotype. Recovery of independent alleles of the same gene rule out a linked functional polymorphism at a different gene as the cause of mutant phenotype. One way to do this with dominant or semi-dominant mutant is to identify loss of function derivatives of the dominant mutant allele. We carried out a suppressor screen to identify loss-of-function alleles of *Sdw2-N1991* to permit molecular identification of the underlying gene. Pollen from *Sdw2-N1991* homozygotes with a severe dwarf phenotype was mutagenized with ethyl methanesulfonate (EMS) and used in controlled pollination of wildtype B73 ears (**Figure 5**). This overwhelmingly produced uniform semi-dwarf *Sdw2-N1991*/+ heterozygous progenies. The rare events of pollen contamination during pollination and EMS-induced mutations that knock out the semi-dominant *Sdw2-N1991* allele, resulted in tall individuals that closely resemble a wildtype B73 plant (**Figure 5**). We screened ∼8,000 M1 plants from this cross and identified eight B73-like plants of normal height. Pollen contamination of our M1 population affected by pollen from neighboring B73 parents, self-contamination of the ear parent by incomplete detasseling, or gynogenetic haploidy could all result in a wildtype B73 plant and would interfere with the identification of true suppressors (**Figure 5**). However, such contaminants are expected to have B73 alleles at the location of *Sdw2-N1991*. Contamination of our crosses with pollen from a non-B73 genotype would result in alternate alleles across the maize genome, and a tall hybrid phenotype. We culled (aka rogued, in the maize breeder’s parlance) three tall hybrids from our population during the field season. Loss of function at the *Sdw2-N1991* allele, however, should result in intragenic suppressor alleles that would retain the alternative alleles encoded at the introgression on chr3 from 15-130Mb identified above and exhibit B73 alleles everywhere else (**Figure 5**).

**Figure 5.**
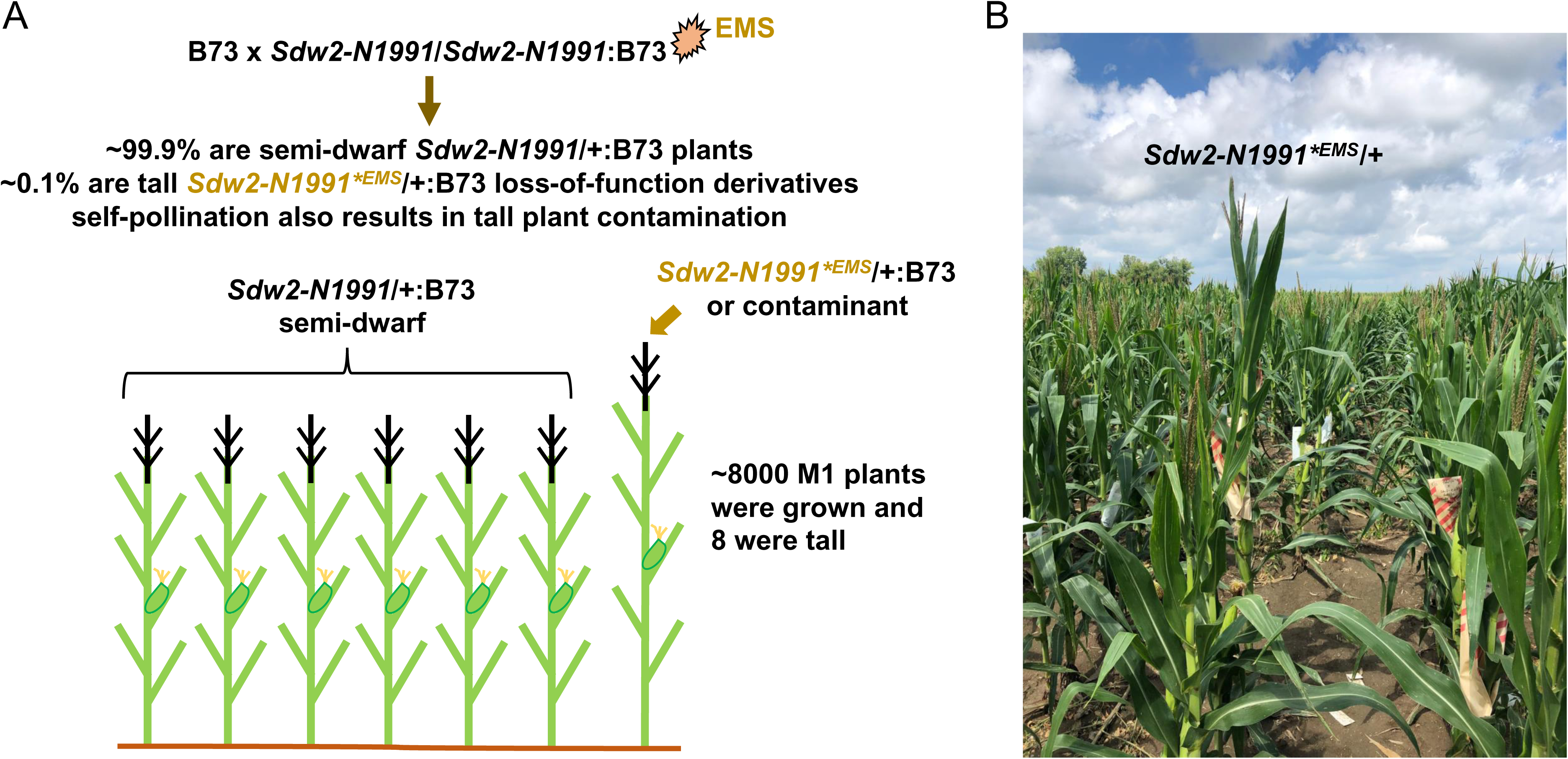
Identification of true suppressors from contaminants in maize pollen EMS mutagenesis. (A) Design of the suppressor screen for loss-of-function derivative alleles of the semi-dominant *Sdw2-N1991* allele using pollen EMS treatment. (B) Observed loss-of-function derivative of *Sdw2-N1991* allele (tall plant in a field of semi-dwarf siblings) in the M1 population from B73 x *Sdw2-N1991*/ *Sdw2-N1991* (pollen EMS).

To detect pollen-contaminants and find true intragenic suppressors of *Sdw2-N1991* allele, we used the WideSeq approach to genotype all eight wildtype B73-like suppressors in the M1 population. The depth of sequencing of these eight samples ranged from 0.006-0.01x (**Table 1**). Paired-end reads were mapped to the B73 v4 reference genome. Sequences overlapping the polymorphic positions in maize (HapMap v3) were tabulated as reference or non-reference and the frequency of non-reference reads in each 5Mb bin was calculated. Seven of the eight B73-like tall plants in our suppressor screen encoded B73 alleles across the entire genome, including at the position of *Sdw2-N1991*, indicating that they are B73 pollen contaminants in our M1 population (**Figure 6**). One tall individual, suppressor 8, had an introgression of non-B73 DNA, as indicated by high non-reference reads frequencies in multiple consecutive bins starting at the 15Mb bin through the bin ending at 130Mb on chr3 at the location of *Sdw2-N1991* (**Figure 6**). These correspond to the window identified for *Sdw2-N1991* shown in **Figure 5**. Thus, suppressor 8 encodes a revertant allele of *Sdw2-N1991* and is designated as *Sdw2-N1991^suppressor8^.* This demonstrates that our economical sequencing approach can be used in combination with EMS mutagenesis suppressor screens to accelerate gene discovery.

**Figure 6.**
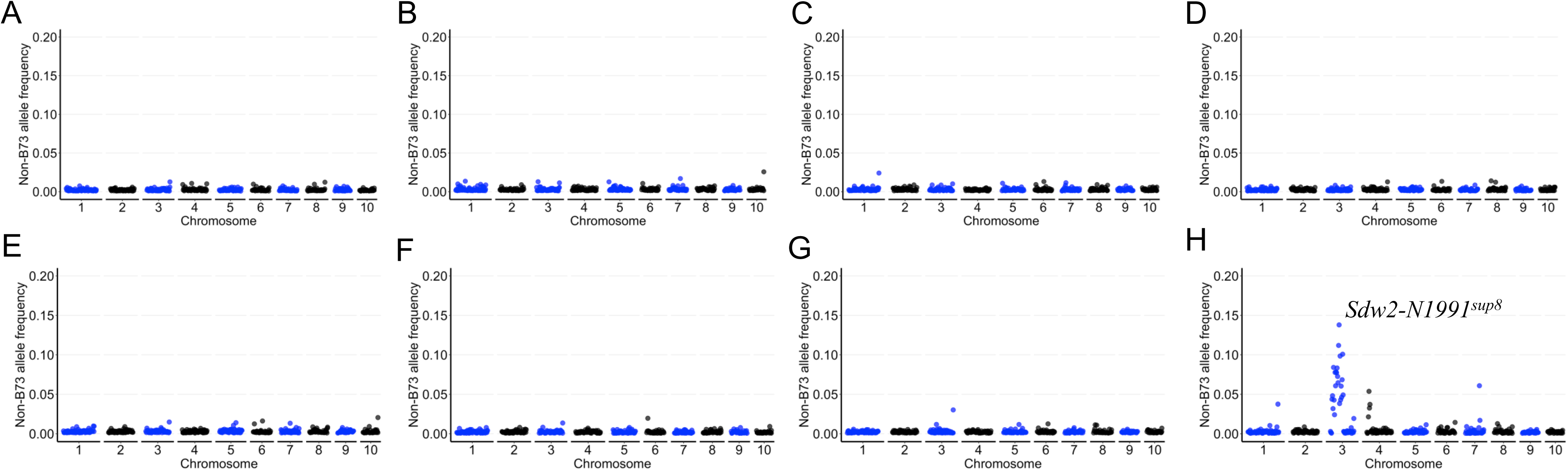
WideSeq genotyping of putative *Sdw2-N1991* derived suppressors. The genome-wide plot of non-B73 allele frequency at 5 Mb bin resolution using the HapMap3 positions across all ten chromosomes of maize in eight tall individuals from B73 x *Sdw2-N1991/Sdw2-N1991* (pollen-EMS) M1 population. (A-G) Suppressors due to self-pollination contaminants of B73 ear parents contain no *Sdw2* introgression, and (H) a true suppressor (*Sdw2-N1991^sup8^*) encoding the *Sdw2-N1991* introgression on chromosome 3. The start and end position of the peak 5Mb bin on chromosome 3 in *Sdw2-N1991^sup8^* (panel H) is annotated.

### Sequence-anchored pseudomarkers for RIL genotyping at 21 USD per sample

We tested whether the low pass sequencing can accurately detect genomic introgressions and predict crossovers. A Nested Maize Association (NAM) RIL, accession Z004E0189 (Buckler et al. 2009) was sequenced at an average depth of 0.009x (**Table 1**). This RIL is derived from the B73 x CML247 cross through single seed decent (Buckler et al. 2009) and was previously genotyped (Elshire et al. 2011; Olukolu et al. 2014). The polymorphic SNPs between CML247 and B73 in the HapMap3 dataset were used to calculate the non-B73 allele frequency at 5Mb resolution in this line (**Figure 7; Tables S4 and S5**). The allele frequencies calculated using all the reads spanning the CML247 polymorphic SNPs traced the previously detected recombination breakpoints in this RIL (**Figure 7 and Tables S4**). Such 5 Mb bins represent a collective genotype likelihood for a genomic region, where the most likely genotype (B73 or CML247) can be used directly in QTL mapping experiments. This binned genotype is similar to the previously proposed idea of pseudo-markers (Sen and Churchill 2001) and results in every line being scored for every genotyped-bin.

**Figure 7.**
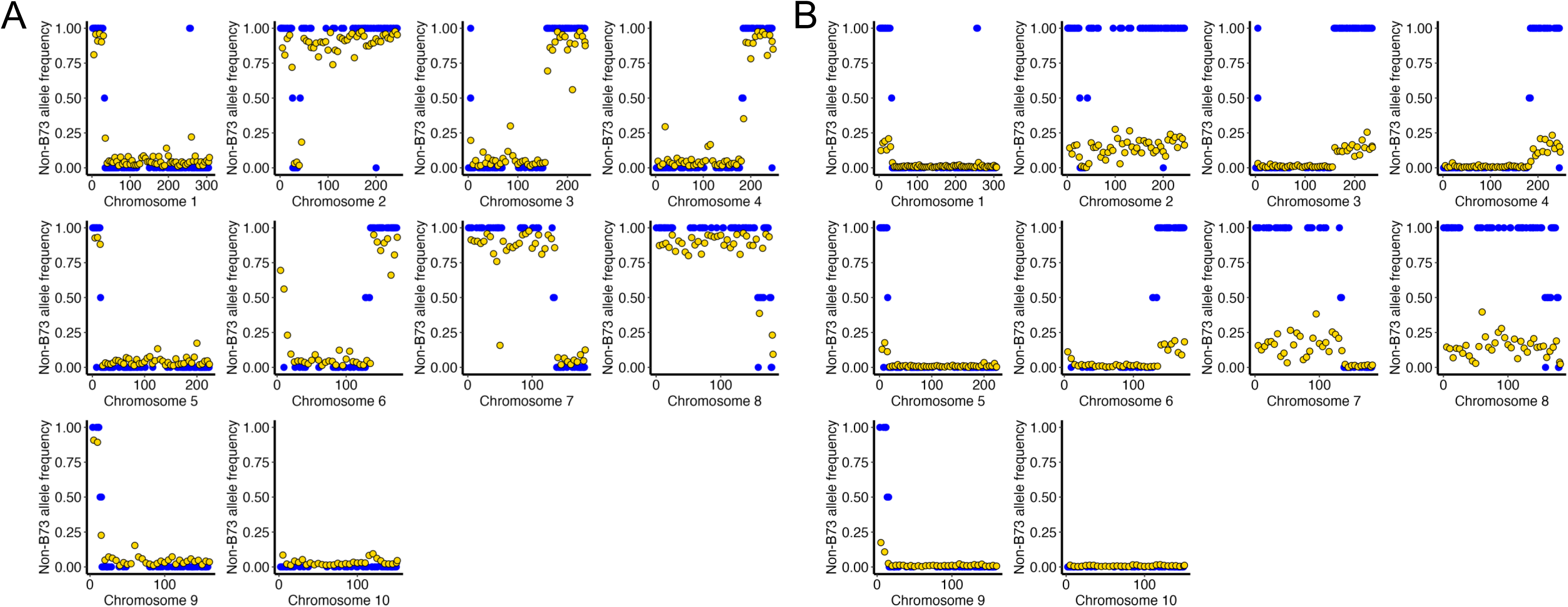
Overlay plot of WideSeq-derived non-B73 allele frequency at 5Mb resolution (golden circles) in the NAM-RIL Z004E0189 and its genotype using 1022 SNP markers (blue) described previously by McMullen *et al*. 2009. The WideSeq non-B73 allele frequency at 5Mb resolution using reads overlapping (A) only CML247/B73 polymorphic SNPs identified in HapMap3, and (B) all HapMap3 SNPs.

To determine if our prior knowledge of the parental genotypes for the Z004E0189 RIL was necessary, we repeated our analysis using all the HapMap3 SNP positions. This calculated the allele frequencies of sequence reads overlapping all the HapMap3 SNPs, instead of only CML247 polymorphic positions. The alternate allele frequencies at the non-B73 genomic regions were clearly visible across the genome and tracked the previous genotypes of this RIL (**Figure 7 and Table S5**). For instance, on chr3 the region spanning from the telomere of the short arm to 155 Mb had an average alternate allele frequency of 0.01, whereas the region from 155 Mb to the end of telomere on the long arm had an average alternate allele frequency of 0.14 (**Figure 7 and Table S5**). A near 14-fold difference in alternate allele frequency makes it easy to discern between the B73 and non-B73 genomic regions. Like we show for *br1-ref*, *br2-ref*, and *Sdw2-N1991*, introgressions into the reference genotype from unknown parents can be easily identified using very little sequence data.

### Determination of introgressed segments and crossovers in Near-Isogenic Lines

We next used near-isogenic lines to test the ability of the low pass sequencing to determine crossover positions and introgressions. We used two NILs from a previously described NIL population where B73 and Mo17 were reciprocally used as recurrent and donor genomes (Eichten et al. 2011). We used the b94 NIL (B73 as the recurrent and Mo17 as the donor parent) and m97 NIL (Mo17 as the recurrent and B73 as the donor parent) to test introgression detection. We previously used b94 and m97 NILs to validate a QTL on chr10 that contained a reciprocal introgressions near the *oil yellow1* (*oy1*) gene at ∼9.2Mb and modify the phenotype to explore expression consequences by RNAseq (Sawers et al. 2006; Khangura et al. 2019; 2020; 2022). We generated $21 of sequencing data using DNA from a single individual of each NIL. The sequence data from b94 and m97 provided 0.01x and 0.005x coverage of the maize genome, respectively (**Table 1**). Any 5Mb bin that was covered by fewer than five reads was excluded from this analysis. All reads overlapping alleles polymorphic between B73 and Mo17 were counted and the number of B73-like and Mo17-like reads were tabulated for each bin. The *oy1* allele in the b94 NIL corresponded to a region that only carried reads of the Mo17 genotype. The homozygous introgression containing the *oy1* locus (∼9.2Mb) extended across the majority of chr10 encompassing bins starting at 5 and continuing to at least 135Mb (**Figure 8 and Table S6**). The b94 NIL was also heterozygous for chr5 starting at 30Mb through 165 Mb and homozygous for Mo17 genotypes at each edge this block. The crossovers and sections of heterozygosity detected by sequencing largely correspond to the recombination breakpoints and allelic states previously determined using a SNP chip (**Figure 8A**; Eichten et al. 2011). To try and find the edge of the introgression from Mo17 on chr10, we recalculated SNP frequencies using 1Mb bins. Some bins contained fewer than five reads and were not analyzed further. The bin from 9-10Mb on chr10 that contains *oy1*, had 32 reads across polymorphic SNPs, and all these reads were Mo17 type (**Table S7**). Subsequent 1Mb bins with more than five reads were also populated with Mo17 alleles, as was also clear with the 5Mb bins (**Figure 8, Tables S6 and S7**). The analysis of non-reference reads in bins towards the telomere from *oy1* indicate a crossover to be somewhere between 5Mb and 7Mb, most likely towards the telomere from 6Mb. The 5-6Mb bin contains 20 reference B73 reads and 1 Mo17 type read and the bins from 6-7Mb, 7-8Mb, 8-9Mb have 3, 55, and 12 reads, respectively, all of Mo17 type (**Table S7**). This prediction matches the genotypic data from Eichten et al. 2011, where the last marker with B73 genotype was on chromosome 10 at 5853032 bp and the first marker with homozygous Mo17 genotype was at 6071650 bp (**Tables S8**). Given the certainty of this crossover region, we looked at each read to identify the narrowest possible interval. The last B73-like read overlapping a B73/Mo17 polymorphic SNP was at 5629603 bp (SNP genotype: T/G) and the first Mo17-like read at 6012706 bp (Indel genotype: -/C), followed by another Mo17-like read at 6747859 bp (SNP genotype: G/A). This maps the recombination between 5629603 and 6012706. If we take the two data types together, the recombination interval is narrowed down between 5853032 and 6012706 on chr10.

**Figure 8.**
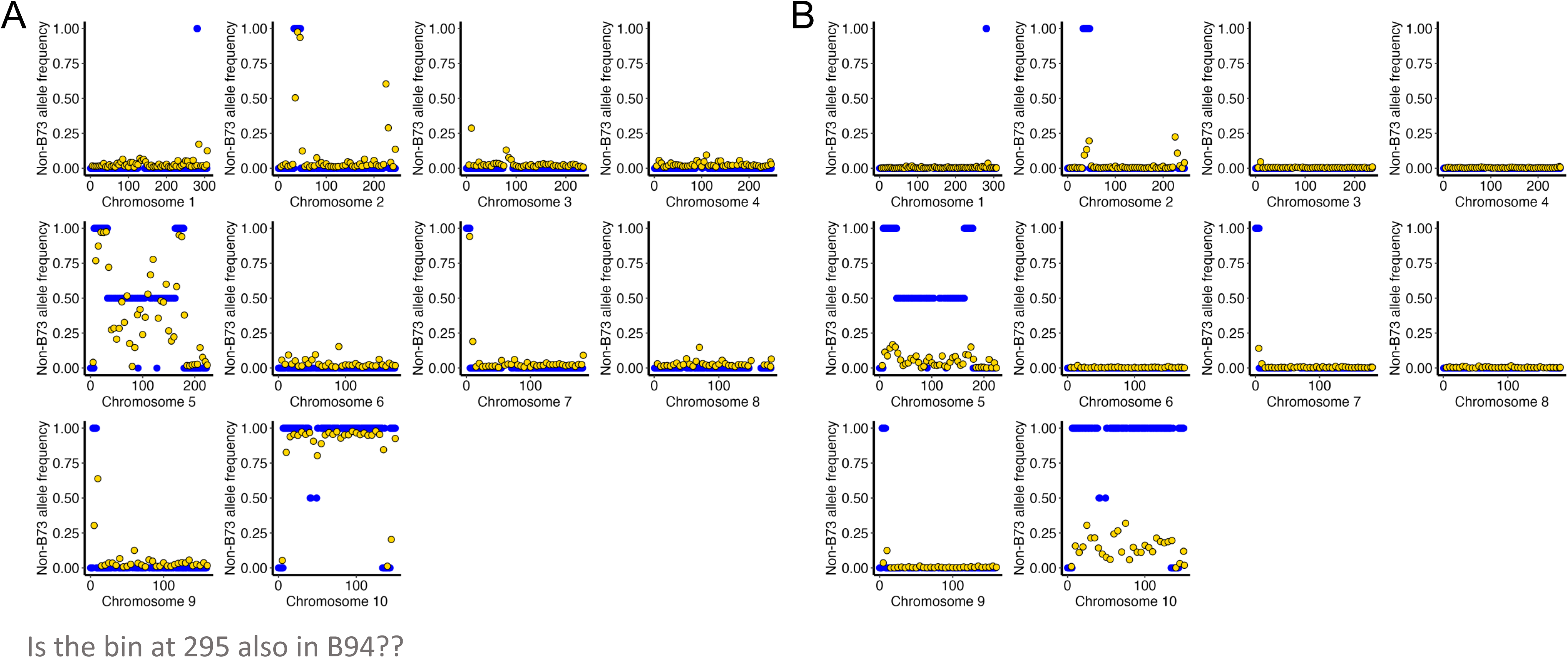
Overlay plot of WideSeq-derived non-B73 allele frequency at 5Mb resolution (golden circles) in the b94 near-isogenic line and its genotype using 50K SNP array (blue circles) described previously by Eichten *et al*. 2011. The non-B73 allele frequency at 5Mb resolution from WideSeq reads in b94 is plotted using reads overlapping (A) only Mo17/B73 polymorphic SNPs identified in HapMap3, and (B) all HapMap3 SNPs.

The m97 NIL was mostly Mo17 with a handful of B73 introgressions on chromosomes 3, 4, 8, and 10 (**Figure 9 and Table S9**). This NIL carries a B73 segment at the top of chr10, from the telomere to ∼10Mb, which includes the *oy1* locus. To estimate the recombination breakpoint more closely, allele frequencies were calculated for 1Mb bins. A close recombinant to the *oy1* locus (∼9.2Mb) is present in m97 as the bin containing *oy1* from 9-10 Mb returned 8 B73-like reads and no Mo17-like reads while the next bin towards the centromere at 10-11 Mb had no B73-like reads and 4 Mo17 reads (**Table S10**). The reads were evaluated across the 9-11 Mb region. This detected the last B73-like read overlapping a B73/Mo17 polymorphic SNP at 9817244 bp (SNP genotype: A/G) and the first Mo17-like read at 10505907 bp (SNP genotype: A/T) and maps the recombination between these two locations. The previous SNP genotyping of m97 NIL identified a homozygous B73 genotype at 9851961 bp and a homozygous Mo17 genotype at 10160759 bp on chromosome 10 (**Table S8;** Eichten et al. 2011). The higher density SNP data provides a closer estimate for the breakpoint, but the correspondence of these windows demonstrates that the low coverage genotypes can be used for crossover detection without the need to produce a SNP array, oligos for sequence-capture, PCR primers at polymorphic positions, or other project-specific reagents.

**Figure 9.**
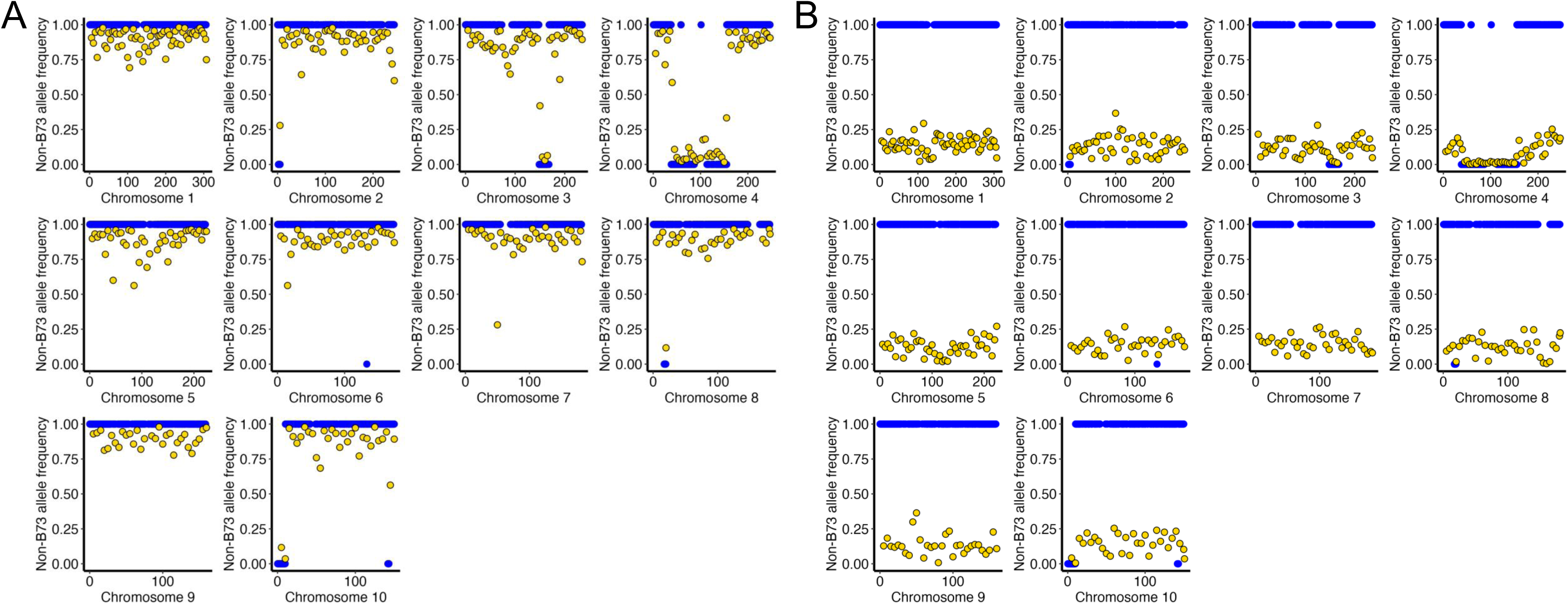
Overlay plot of WideSeq-derived non-B73 allele frequency at 5Mb resolution (golden circles) in the m97 near-isogenic line and its genotype using microarray probes (blue circles) described previously by Eichten *et al*. 2011. The non-B73 allele frequency at 5Mb resolution from WideSeq reads in m97 is plotted using reads overlapping (A) only Mo17/B73 polymorphic SNPs identified in HapMap3, and (B) all HapMap3 SNPs.

To demonstrate the capability of this approach to detect non-B73 introgressions we repeated analysis using all the HapMap3 SNPs. The Mo17 introgression on chr10 from 55-130 Mb in the b94 NIL had an average alternate allele frequency of 0.17 when all HapMap3 SNPs were tallied (**Figure 8 and Table S11**). In m97, the alternate allele frequency at all the Mo17 introgressions on chr1, chr5, chr7, and chr9 that had no B73 introgression averaged around 0.14, 0.12, 0.15, and 0.13 (**Figure 9 and Table S12**). In contrast the B73 introgression on chr4 from 45-150 Mb had average alternate allele frequency of 0.01. This is consistent with our ability to detect *br2-ref*, *br1-ref*, and *Sdw2-N1991* introgressions using the HapMap3 SNPs demonstrated above (**Figures 2, 3, and 4**).

### Forensic haplotype identification of known introgressions using low-pass sequencing data

The abundance of non-B73 SNPs in our reads suggested that we might be able to identify the most similar donor haplotype for each of our unknown mutants in the HapMap3 dataset that contains 1210 maize accessions (Bukowski et al. 2018). To test the feasibility of this, we first used data from the b94 NIL and Z004E0189 RIL, to see if this correctly identified the known parental line. The b94 NIL carries Mo17 introgressions in an otherwise B73 background (Eichten et al. 2011). The genotypes at all HapMap 3 SNPs across the 9-19Mb window encoding the introgression of *oy1* were tabulated (Table S13). There were 168 non-B73 SNPs between 9-19Mb on chr10 in b94. Four SNPs with MAF below 0.05 in HapMap3 were eliminated to reduce the influence of error prone positions. The remaining 164 SNPs were compared to the genotypes of the 1210 accessions present in the maize HapMap3 panel. We calculated the genotypic similarity, quantified as a Jaccard Index (Gilbert 1884; Jaccard 1901), of each HapMap3 accession with the b94 NIL introgression. This region showed highest genetic similarity of 98.2% with LH52, followed by second highest similarity of 97.6% with Mo17 (**Figure 10** and **Table S13**). Similarly, the other sample Z004E0189 NAM-RIL is derived from a cross between CML247 and B73. We used a small genomic region to carry out a haplotype analysis from this line. We chose 80-85Mb on chr8 which has an elevated non-B73 allele frequency using HapMap3 SNPs (**Figure 10**; **Tables S14**) and was previously identified as homozygous for the CML247 genotype (McMullen et al. 2009). We identified 225 non-B73 SNPs in this window in Z004E0189. Genetic similarity was calculated (Gilbert 1884; Jaccard 1901) by comparing the genotypes from Z004E0189 to the 1210 accessions in the maize HapMap3 panel (**Table S14**). The highest genetic similarity of 98.7% was returned for CML247. The high accuracy of haplotype prediction using limited sequence data in both b94 NIL and Z004E0189 RIL demonstrates that low-pass sequencing is sufficient to assess pedigree-of-origin in maize. Our ability to assess parent-of-origin in these known pedigreed materials may allow researchers to determine the origins of introgressions in material that lacks pedigree information by comparing low pass genotypes to the set of haplotypes in the haplotype map of their species of interest.

**Figure 10.**
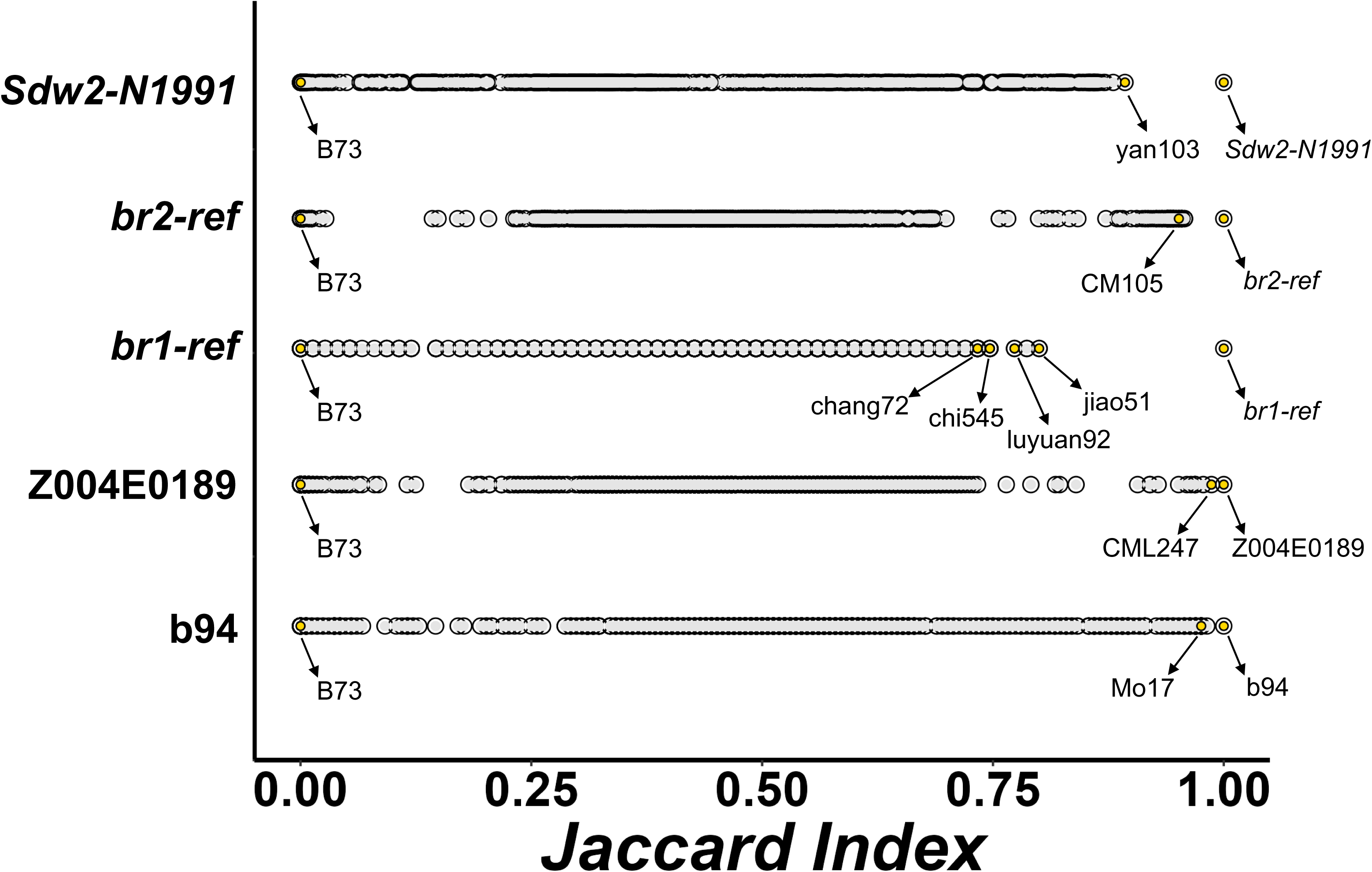
Genetic similarity (quantified as Jaccard Index on the x-axis, using only non-B73 reference calls) in the test sample (displayed on the y-axis) against the 1210 maize HapMap3 accessions. The genomic regions tested using WideSeq are as follows: b94 (chr10:9-19Mb); Z004E0189 (chr8:80-95Mb); *br1-ref* (chr1:215-220Mb); *br2-ref* (chr1:202-210Mb); *Sdw2-N1991* (chr3:55-110Mb).

**Figure 11.**
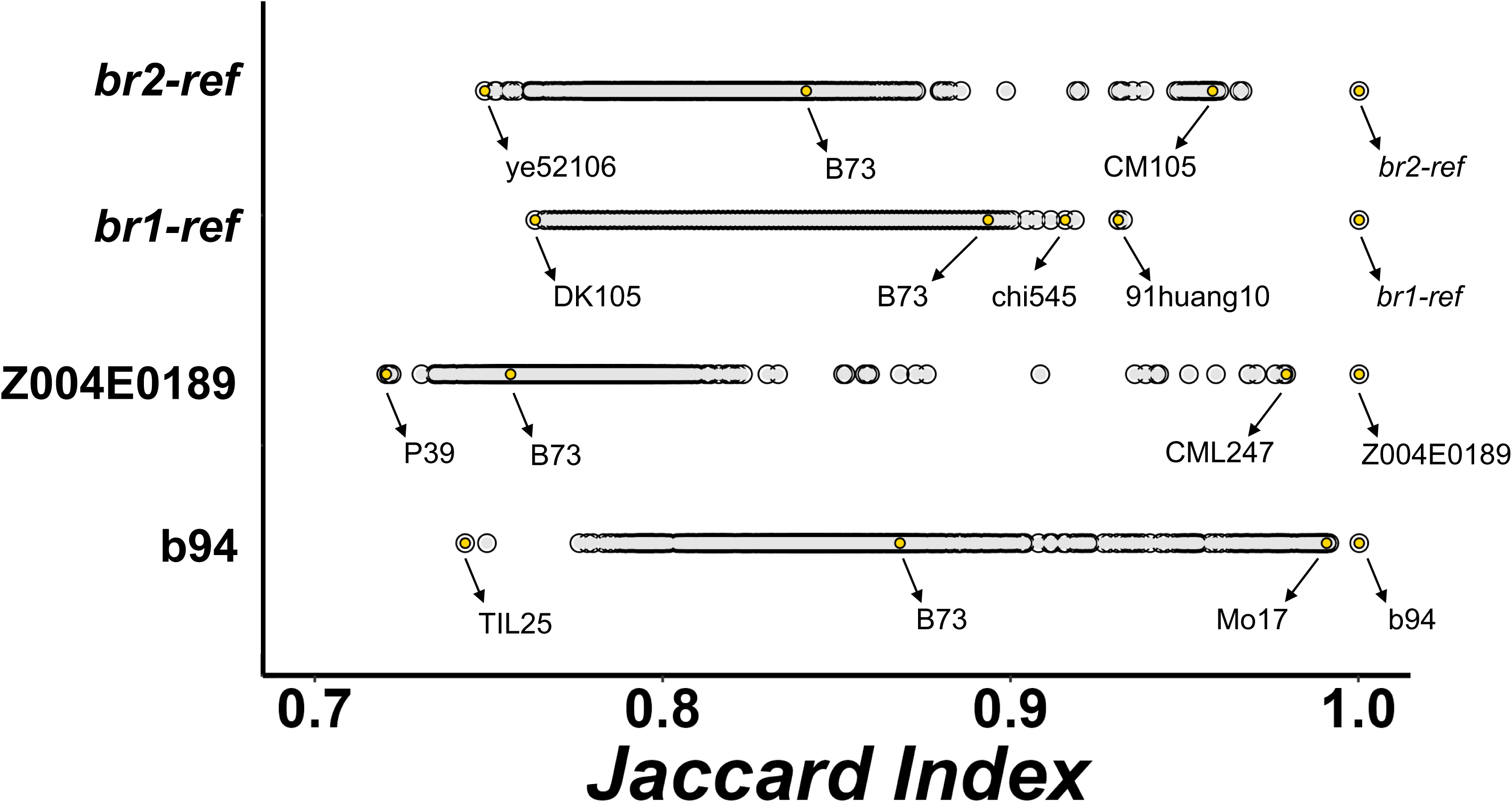
Genetic similarity (quantified as Jaccard Index on the x-axis, using both alternate and reference positions) in the test sample (displayed on the y-axis) against the 1210 maize HapMap3 accessions. The genomic regions tested using WideSeq are as follows: b94 (chr10:9-19Mb); Z004E0189 (chr8:80-95Mb); *br1-ref* (chr1:215-220Mb); *br2-ref* (chr1:202-210Mb).

We tested whether the haplotype analysis presented here using low pass sequencing could be used to detect pedigree information from RNAseq data as well. SNPs were called after aligning reads from RNAseq data from b94 x B73 F1 hybrids seedlings to the B73 genome (Kaur et al. 2025). SNPs from the same 9-19Mb region used above were tabulated. Using all the HapMap3 SNPs in this window, we identified 213 high-quality diagnostic SNPs that were heterozygous in the b94 x B73 F1 hybrids in this region (**Table S15**). Thirty-one maize accessions, including Mo17, showed 100% genetic similarity with the b94 sample. Therefore, high quality SNP data from RNAseq can predict the haplotype with extremely high accuracy in the b94 x B73 F1 hybrid seedlings. That said, when a haplotype is common among the genotyped lines, as is the case here for Mo17 (and see below for inbred B14), this may not identify a single line as the most likely progenitor.

### Haplotype estimation of mutant allele donors from lines with incomplete pedigree information validates introgressions and historical records

The lack of information on the origin of genetic stocks can complicate genetic studies aimed at understanding the molecular change in the mutant allele that is associated with a phenotype. Given our successful inference of pedigree for the b94 NIL and the Z004E0189 NAM-RIL demonstrated above, we tested the ability of our pipeline to predict the parent of the mutant alleles described in this study that have incomplete pedigree records. We used the non-B73 introgressions where the mutant alleles *br1-ref*, *br2-ref*, and *Sdw2-N1991* map to identify maize inbred lines from HapMap3 with highest genetic similarity.

The homozygous *br1-ref* pool yielded the highest non-B73 allele frequency from 215-220Mb on chr1, and the *br1* gene itself resides at ∼216Mb (**Figure 3**). Interrogating all the non-reference SNP calls in this genomic interval, we identified 75 diagnostic non-B73 SNPs in the *br1-ref* sample. Multiple accessions from Chinese breeding programs displayed high genetic similarity at these 75 SNPs with jiao51 having the closest match (**Figure 10 and Table S16**). The *br1-ref* allele was identified as a spontaneous mutant segregating within a *waxy* maize population from China that was hybridized to Algerian germplasm, and later introgressed into North American accessions (Kempton 1921). USDA-ARS Maize Genetics COOP aquired this line from Prof. Rollins A. Emerson in 1943 (crosses Em. 28-608 (6)/Em 28-44 (9)) who likely obtained it from James Howard Kempton (M. Sachs personal communication). The large number of Chinese maize accessions with high genetic similarity with *br1-ref* allele provides a molecular corroboration with the historical record that the reference allele *br1-ref* used in our work arose on the Chinese haplotype and is the same allele described by Kempton in 1921.

We also carried out haplotype prediction using the reads on either side of the *br2-ref* allele. The parentage of this allele is complicated by the incomplete record of 20^th^ century maize breeding. Multiple *br2* alleles may have arisen at different times. Singleton refers to a *br2* allele as derived from the R4 inbred (Singleton 1959). Other alleles were reported as arising spontaneously in the CM104 inbred line, one of a number of lines produced at the Manitoba field station in Canada that along with CM105 have been used in a number of breeding programs (Brewbaker 2011; Troyer 2004; Garg et al. 2007). These lines are inbred lines derived from backcrosses of maize line V3 to the B14 inbred (Reid et al. 2011; Garg et al. 2007; Gethi et al. 2002). The stock we used for our experiments is a double mutant for *hm1* (∼202Mb) and *br2-ref* (∼204Mb) and may have been constructed by Paul Sisco in 1989 (Marty Sachs, personal communication). If this is true, then the *br2* allele should be flanked by DNA derived from CM104, and a recombination should be evident between the *br2-ref* allele and the haplotype harboring the *hm1* mutant allele. The CM104 accession is not part of the HapMap3 panel, but CM104 shares high genetic similarity with the related inbred line CM105 (Garg et al. 2007), which is in the HapMap3 panel. We used the region from 202Mb to 210Mb to predict the haplotype surrounding *br2-ref* (**Table S17**). This identified 329 diagnostic non-reference HapMap3 SNPs. The inbred CM105 was identical to *br2-ref* at 95.1% of these 329 SNPs (**Figure 10 and Table S17**). 64 inbred lines had 95% or better identity with the 8 Mb region on chr1 between 202-210 Mb that contains *br2-ref* (**Table S17**). Of those that had publicly available pedigree information (https://github.com/RILAB/historical_genomics), most lines had B14 as a major component of their pedigree. This indicates that the *br2-ref* allele used in the construction of the *hm1 br2* double mutant and *hm1 hm2 br2* triple mutant stocks (stocks 114F and 114G) at the USDA-ARS Maize Genetics COOP is the allele that arose in CM104 described previously (Brewbaker 2011). It also demonstrates that we can detect the haplotype and origin of classical maize mutants utilizing a combination of historical pedigree information and the available population-wide genetic polymorphisms described in the HapMap. The allele at *hm1* in this material contains an insertion that disrupts the 4^th^ exon of the gene (Multani et al., 1998). The haplotype of this *hm1* mutant allele is unknown. We tested the SNPs closest to *hm1* and found no clear match within the HapMap3.

The *Sdw2-N1991* mutant allele was generated and donated to the Maize Genetics COOP by the late Dr. Gerry Neuffer. At the Maize Genetics COOP, stocks carrying the *Sdw2-N1991* allele lack complete records of the genetic background used for mutagenesis and propagation. To identify the closest-matching maize lines or a likely donor genome for the *Sdw2-N1991* allele, we performed haplotype detection as described previously. We identified 4383 non-B73 HapMap3 SNP positions in the genomic region spanning 55-110 Mb (**Table S20**), which includes the *sdw2* gene (**Figure 3**). No inbred lines favored by Dr. Neuffer were among the top SNP matches for this mutant (**Figure 10 and Table S20**). This suggests that the maize accession mutagenized to generate *Sdw2-N1991* is not present in the HapMap3 set or that a recombination between haplotypes occurred during the maintenance of this stock.

## Discussion

This study details the utility of a skim sequencing strategy, called WideSeq at Purdue, for a variety of genome-wide genetic analyses. The ease of library preparation using Tn5-based tagmentation makes it easy to adapt with minimal additional investment in project-specific reagents. WideSeq was developed by the Purdue Genomics Facility as a service to sequence PCR pools and *de novo* assemble PCR amplicons, plasmids, and BACs. We demonstrated the utility of very low-pass sequencing for genome-wide genetic analyses in maize and other species, provided that both a high-quality reference genome and polymorphic sites are already known. By combining a small number of reads with known SNP positions, even in the large and complex genome of maize, this method can map genes, identify introgressions, screen contaminants in a population, identify crossovers in near-isogenic and recombinant inbred lines, and identify the haplotype-of-origin materials with unknown parentage. This approach worked even at sequencing depths as low as 0.014x (**Table 1)**.

Our mapping experiments were enabled by 55 million HapMap3 SNP polymorphisms in maize, determined from a collection of 1210 maize accessions (Bukowski et al. 2018). The low sequencing depth limited our ability to call HapMap3 SNP positions as homozygous or heterozygous. Therefore, we calculated allele frequencies over contiguous 5Mb genomic intervals. To do this, each sequence that overlapped a HapMap3 SNP was tabulated as either encoding the reference or non-reference SNP allele for all reads in each bin (**Figure 1**). The inclusion of both reference and non-reference positions for haplotype calling provided a clear signal of haplotype match in homozygous samples. In heterozygotes, these calculations are more complicated and must rely on the non-reference reads. In an introgressed segment in a heterozygote, even in a recurrent cross to the reference genotype, all non-reference reads at SNP positions are unequivocally interpreted as derived from the presence of a non-reference genotype in the DNA sample. Any read that contains the reference genotype either corresponds to the recurrent parent contribution in the heterozygote, or because the haplotype of the introgression donor matches the reference type.

As a proof-of-concept, we first determined the introgressions of two recessive mutant alleles *br1-ref* and *br2-ref*, that have been molecularly characterized (**Figures 2 and 3**; Multani et al. 2003; Jiao et al. 2023). Even in the absence of any known parent-specific polymorphisms for *br1-ref* and *br2-ref* mutants, we successfully mapped the genomic introgressions of both alleles using the HapMap3 SNPs positions. We next mapped an uncharacterized dominant mutant *Sdw2-N1991* using DNA pools from heterozygous mutants and wildtype siblings from an advanced backcross population (**Figure 4A**). The parentage of *Sdw2-N1991* is unknown. Using HapMap3 SNPs, we identified a non-B73 introgression on chromosome 3 from 15-130 Mb that contains the *sdw2* locus (**Figure 4B**). This allowed us to efficiently remove contaminants from an EMS suppressor screen to knock out the dominant *Sdw2-N1991* allele (**Figure 5**). This suppressor screen contained eight plants that phenocopied wildtype B73 plants. Seven of the eight putative suppressors were eliminated as B73 contaminants using $21 of sequencing. This also identified a true suppressor allele, *Sdw2-N1991^suppressor8^*, as indicated by the non-B73 introgression on chromosome 3 (**Figure 6**). This demonstrates that the genome-enabled low-pass genotyping in maize can be used to identify true positives in EMS suppressor screens and can facilitate gene discovery.

This same approach can be used to genotype near-isogenic lines, recombinant inbred lines, and identify recombination breakpoints. The previously generated genotypic data on B73-Mo17 NILs (Eichten et al. 2011) and the NAM-RIL (McMullen et al. 2009) corresponded with the genotypes derived using our low-pass sequencing and bioinformatics pipeline (**Figures 7, 8, and 9**). The low-pass sequence data in the b94 NIL that carries the Mo17 donor introgressions and Z004E0189 NAM-RIL (B73 x CML247) allowed us to test our ability to predict the background-of-origin using the low-pass sequence data. Using reads overlapping multiple HapMap3 SNP in a non-B73 genomic region in both samples, we successfully identified Mo17 among the top match in b94 NIL, and CML247 in the Z004E0189 NAM-RIL (**Figure 10**). This demonstrates that we can use low-pass sequencing and known polymorphisms to identify the background of origin for mutants of unknown parentage.

The non-B73 introgressions in the *br1-ref* and *br2-ref* mutants were also used to determine the likely parent-of-origin for the haplotypes surrounding these alleles. Using the genetic similarity of the informative SNPs at the introgression flanking these two genes, we identified inbred lines in the HapMap3 panel that carry related haplotypes (**Figure 10**). Literature searches for historical records of the background of both mutant alleles provide some corroboration, and as such, we provide the first molecular insight into the likely genetic backgrounds of the alleles *br1-ref*, from a Chinese variety, and *br2-ref*, from a Canadian line derived from B14. Remarkably, in 1921, Kempton speculated, based on the lack of other brachytic plants in his nursery, that a single gamete from a Chinese open pollinated variety gave rise to the novel brachytic mutant (Kempton 1921), and we can now confirm that this speculation was correct as the *br1-ref* allele occurs within the Chinese haplotype. We believe similar haplotype analysis could be done across the hundreds of mutant stocks at the Maize Genetics COOP that were collected as spontaneous mutants of unknown pedigree or for which pedigree information has been lost. This would facilitate gene discovery by allowing researchers to map these mutants using the SNPs in HapMap3 from known haplotypes. Furthermore, this approach will work in any species with known variants, including mouse, Arabidopsis, crops, and livestock. Given the extreme pressures on genetic stock centers, such an approach would permit the inexpensive detection of stock duplications to reduce genetic redundancy in seed collections and improve the sustainability of these resources.

### Ultra-low coverage sequencing of large genomes works when they are diverse

The success of such low sequencing depths in maize demonstrates the value of considering genomic windows, recombination, SNP diversity rates, and gene numbers when planning genomic experiments. For species with large genomes, there is often an expectation that higher data throughput will be required for genomic approaches to work. However, multiple considerations can reduce the coverage required to conduct genetic analyses such as the haplotyping and mapping methods we presented here. For example, the number of reads overlapping any SNP determines the rate at which reads are informative. A smaller genome with fewer polymorphisms between haplotypes (e.g., Arabidopsis) will produce a higher proportion of uninformative reads and thus require a *higher* sequencing depth to resolve recombination or detect the origin of haplotypes than was required for the highly polymorphic maize material used here. If we were working with lower SNP rates (e.g., EMS mutants) the same coverage would be needed for both Arabidopsis and Maize, and much more data would be required to reach that coverage in a species with 2.5 Gbp per chromosome complement than 155 Mbp. Similarly, when working with RILs, NILs, or line crosses, the greater the recombinational length of the genome the more reads that are required to resolve all the recombinants. Recombinational length is dramatically altered by chromosome number, and two genomes of the same size in Mb but differ two-fold in chromosome number while require different numbers of reads to provide the same resolution *per chromosome*. Similarly, if the goal is to identify windows that contain a set number of genes (e.g. map a locus to a window with a single gene) then the recombination length of the genome expressed as a per-gene dimension determines both the size of the population required and the depth of sequencing necessary to achieve this. Similar effects occur in GWAS experiments and in determining the depth required to accurate impute SNP data across a population. A combination of the SNP density, kinships, and the LD decay in the genome will determine how much sequencing depth is required to accurately determine the genotype of a line. These values, rather than solely relying on the size of the genome, respond to genetic and population genetic parameters. Inclusion of this in the design of experiments can both increase the success of mapping and greatly increase the efficiency of experimentation by preventing researchers from sequencing at depths responsive to the length of the genome rather than its diversity.

### If you want to go far, go together: data sharing enables genomics at a lower price

Altogether, these analyses demonstrate that mutants of unknown origin can be mapped using population level SNP genotypes, such as the maize HapMap, using very low sequencing coverage. This untargeted low-pass sequencing can carry out haplotype predictions. We were able to achieve linkage mapping and haplotype prediction using very little sequencing data that cost us a fraction of a typical resequencing experiment in maize. This approach can be easily adapted to any organism so long as a haplotype map exists. In this sense, our ability to do experiments at this low price point represents a “post-genomic” approach. This study was facilitated by large prior public investment in sequencing, genome analysis, as well as the public data sharing through publicly funded data repositories built by the maize genetics community, the USDA, and the NSF. We understand that not all plant species currently have similar resources. A collective effort to place genomic data in the public domain for unrestricted use, could trigger development of similar resources for other plant species. Through a wide range of public datasets such as resequencing and transcriptomic datasets, haplotype maps could be built. This would allow use of population-level SNPs to perform tasks like linkage mapping. Over time as more resequencing datasets accumulate, haplotype maps could be enriched to capture even more population level SNPs. Therefore, we encourage similar collaborations in other organisms to enable and accelerate scientific discoveries.

Introgressions were easiest to detect when the reference genome used for alignment and for the HapMap corresponded to one of the parents. For maize, this genotype is the B73 line. When introgressions were compared to the reference genome the relative change in allele frequency can be very high, even without knowing the correct haplotype. For example, deliberately ignoring the background of the b94 NIL still detected the introgressed segment (**Figure 6B**). The average HapMap3 non-reference allele frequencies on chromosomes 3, 4, 6, and 8, that carried no Mo17 introgressions, were 0.006, 0.004, 0.005, 0.005 and this jumped to 30-fold greater non-reference allele frequencies at the known Mo17 introgressions. Similarly, non-reference alleles were visible in the same locations of independent pedigrees of B73 backcrosses for *br1*, *br2*, and *sdw2*. The provenance of the B73 that these mutants were introgressed into was unknown. Polymorphisms between B73 sources was found in a comparison of multiple transcriptomic experiments done across 20 research groups using B73 with an average of 2.3% unintentionally fixed polymorphisms (Liang and Schnable 2016). We have since ordered the reference genotype from the USDA-ARS GRIN (PI 550473) and confirmed that PI 550473 corresponds to the sequenced B73 and does not have these non-B73 introgressions.

### Low-pass genotyping for efficient use of resources

In addition to the use cases presented in this study, low coverage sequencing approaches can achieve various teaching and research objectives. Genomics can be complex to integrate into undergraduate curricula due to limited access to wet lab resources, complex reagents, data costs, and computational burdens in analysis. Best practices for sequencing data handling, understand the basics of sequence alignment, linkage mapping, and forensic haplotype analyses outlined in this study could be easily taught in the classroom. The low data volumes used here allow genome-wide analyses to be carried out on a standard consumer laptop making it possible to conduct hands-on training for bioinformatics beginners. The ability to identify haplotypes using low-pass data can also benefit breeding programs for trait introgression, foreground and background selection, and rogue any contaminants. The low cost per sample could enable more efficient and rigorous intellectual property protections for breeding germplasm from small public breeding programs, for instance.

## Author Contributions

P.S.M developed and implemented the WideSeq method, R.S.K. and B.P.D. designed research, R.S.K., P.S.M. performed experiments, R.S.K., A.K., and B.P.D. analyzed data, R.S.K. and B.P.D. wrote the manuscript, and all authors read and approved the manuscript.

## Acknowledgments

We want to acknowledge the staff at the Purdue Genomic Facility for developing and implementing the WideSeq protocol. We would also like to thank Jim Beaty and Rachel Stevens at the Purdue Agronomy Center for Research and Education, Jason Adams at the Indiana Corn and Soybean Innovation Center, and Nathan Deppe at the Purdue Horticulture Greenhouse for their help in growing plants and handling plant materials used for experiments in this study. We gratefully acknowledge Ford Prefect, visiting editor, for reading and improving this manuscript by adding an adverb. RSK was supported by the USDA NIFA Postdoctoral Fellowship award# 2022-67012-36601. This work was supported by the National Science Foundation IOS-2309932 to BPD.

## Reference

Addo-Quaye, Charles, Mitch Tuinstra, Nicola Carraro, Clifford Weil, and Brian P. Dilkes. 2018. “Whole-Genome Sequence Accuracy Is Improved by Replication in a Population of Mutagenized Sorghum.” G3 (Bethesda, Md.) (England) 8 (3): 1079–94. 10.1534/g3.117.300301.

Auton, Adam, Gonçalo R. Abecasis, David M. Altshuler, et al. 2015. “A Global Reference for Human Genetic Variation.” Nature 526 (7571): 68–74. 10.1038/nature15393.

Baird, Nathan A., Paul D. Etter, Tressa S. Atwood, et al. 2008. “Rapid SNP Discovery and Genetic Mapping Using Sequenced RAD Markers.” PloS One (United States) 3 (10): e3376. 10.1371/journal.pone.0003376.

Bolger, Anthony M., Marc Lohse, and Bjoern Usadel. 2014. “Trimmomatic: A Flexible Trimmer for Illumina Sequence Data.” Bioinformatics 30 (15): 2114–20. 10.1093/bioinformatics/btu170.

Brewbaker, James L. 2011. “Descriptions of Near-Isogenic Lines of Inbred Hi27.” Maize Genetics Cooperation Newsletter 85: 1–24.

Buckler, Edward S., James B. Holland, Peter J. Bradbury, et al. 2009. “The Genetic Architecture of Maize Flowering Time.” Science (New York, N.Y.) (United States) 325 (5941): 714–18. 10.1126/science.1174276.

Bukowski, Robert, Xiaosen Guo, Yanli Lu, et al. 2018. “Construction of the Third-Generation Zea Mays Haplotype Map.” GigaScience 7 (4). 10.1093/gigascience/gix134.

Burow, Katelin M., Xi Yang, Yun Zhou, Brian P. Dilkes, and Jennifer H. Wisecaver. 2024. “A Brassinosteroid Receptor Kinase Is Required for Sex Determination in the Homosporous Fern Ceratopteris Richardii.” bioRxiv, ahead of print. 10.1101/2024.10.09.617452.

Candela, Héctor, and Sarah Hake. 2008. “The Art and Design of Genetic Screens: Maize.” Nature Reviews. Genetics (England) 9 (3): 192–203. 10.1038/nrg2291.

Danecek, Petr, Adam Auton, Goncalo Abecasis, et al. 2011. “The Variant Call Format and VCFtools.” Bioinformatics 27 (15): 2156–58. 10.1093/bioinformatics/btr330.

Danecek, Petr, James K. Bonfield, Jennifer Liddle, et al. 2021. “Twelve Years of SAMtools and BCFtools.” GigaScience (United States) 10 (2). 10.1093/gigascience/giab008.

Dotto, Marcela C., Jennifer S. Jaqueth, Kevin Fengler, Bailin Li, and Marja C.P. Timmermans. 2010. “Role of Lxm1 in Leaf Patterning and Development.” Poster. 52nd Annual Maize Genetics Meeting, Riva del Garda, Italy.

Eichten, Steven R., Jillian M. Foerster, Natalia de Leon, et al. 2011. “B73-Mo17 Near-Isogenic Lines Demonstrate Dispersed Structural Variation in Maize.” Plant Physiology 156 (4): 1679–90. 10.1104/pp.111.174748.

Elshire, Robert J., Jeffrey C. Glaubitz, Qi Sun, et al. 2011. “A Robust, Simple Genotyping-by-Sequencing (GBS) Approach for High Diversity Species.” PLOS ONE 6 (5): 1–10. 10.1371/journal.pone.0019379.

Garg, Anshu, B. M. Prasanna, and S. V. S. Chauhan. 2007. “Simple Sequence Repeat (SSR) Polymorphism in the Tropical Asian Maize Inbred Lines Differing in Resistance to Banded Leaf and Sheath Blight (Rhizoctonia so/Ani f. Sp. Sasakil).” INDIAN JOURNAL OF GENETICS AND PLANT BREEDING 67 (03): 238–42..

Gethi, James G., Joanne A. Labate, Kendall R. Lamkey, Margaret E. Smith, and Stephen Kresovich. 2002. “SSR Variation in Important U.S. Maize Inbred Lines.” Crop Science 42 (3): 951–57. 10.2135/cropsci2002.9510.

Gilbert, G. K. 1884. “Finley’s Tornado Predictions.” American Meteorological Journal 1: 166–72.

Jaccard, Paul. 1901. “Étude Comparative de La Distribution Florale Dans Une Portion Des Alpes et Des Jura.” Bulletin de La Société Vaudoise Des Sciences Naturelles 37: 547–79.

Jiao, Shuping, Sujan Mamidi, Mark A. Chamberlin, et al. 2023. “Parallel Tuning of Semi-Dwarfism via Differential Splicing of Brachytic1 in Commercial Maize and Smallholder Sorghum.” New Phytologist 240 (5): 1930–43. 10.1111/nph.19273.

Jupe, Florian, Kamil Witek, Walter Verweij, et al. 2013. “Resistance Gene Enrichment Sequencing (R En S Eq) Enables Reannotation of the NB - LRR Gene Family from Sequenced Plant Genomes and Rapid Mapping of Resistance Loci in Segregating Populations.” The Plant Journal 76 (3): 530–44. 10.1111/tpj.12307.

Karre, Shailesh, Saet-Byul Kim, Bong-Suk Kim, et al. 2021. “Maize Plants Chimeric for an Autoactive Resistance Gene Display a Cell-Autonomous Hypersensitive Response but Non-Cell Autonomous Defense Signaling.” Molecular Plant-Microbe Interactions : MPMI (United States) 34 (6): 606–16. 10.1094/MPMI-04-20-0091-R.

Kaur, Amanpreet, Norman B Best, Thomas Hartwig, et al. 2024. “A Maize Semi-Dwarf Mutant Reveals a GRAS Transcription Factor Involved in Brassinosteroid Signaling.” Plant Physiology 195 (4): 3072–96. 10.1093/plphys/kiae147.

Kaur, Amanpreet, Rajdeep S Khangura, and Brian P Dilkes. 2025. “Natural Variation in Tetrapyrrole Biosynthetic Enzymes and Their Regulation Modifies the Maize Chlorophyll Mutant Oy1-N1989.” Plant Physiology 199 (3): kiaf431. 10.1093/plphys/kiaf431.

Kempton, J.H. 1921. “A Brachytic Variation in Maize.” USDA Bulletin 925: 1–28.

Khangura, Rajdeep S., Gurmukh S. Johal, and Brian P. Dilkes. 2022. “Genome-Wide Association Identifies Impacts of Chlorophyll Levels on Reproductive Maturity and Architecture in Maize.” bioRxiv, 2022.11.07.515492. 10.1101/2022.11.07.515492.

Khangura, Rajdeep S, Sandeep Marla, Bala P Venkata, Nicholas J Heller, Gurmukh S Johal, and Brian P Dilkes. 2019. “A Very Oil Yellow1 Modifier of the Oil Yellow1-N1989 Allele Uncovers a Cryptic Phenotypic Impact of Cis-Regulatory Variation in Maize.” G3 Genes|Genomes|Genetics 9 (2): 375–90. 10.1534/g3.118.200798.

Khangura, Rajdeep S., Bala P. Venkata, Sandeep R. Marla, et al. 2020. “Interaction Between Induced and Natural Variation at *Oil Yellow1* Delays Reproductive Maturity in Maize.” G3 Genes|Genomes|Genetics 10 (2): 797–810. 10.1534/g3.119.400838.

Lack, Justin B., Jeremy D. Lange, Alison D. Tang, Russell B. Corbett-Detig, and John E. Pool. 2016. “A Thousand Fly Genomes: An Expanded Drosophila Genome Nexus.” Molecular Biology and Evolution (United States) 33 (12): 3308–13. 10.1093/molbev/msw195.

Li, Heng. 2011. “A Statistical Framework for SNP Calling, Mutation Discovery, Association Mapping and Population Genetical Parameter Estimation from Sequencing Data.” Bioinformatics (Oxford, England) (England) 27 (21): 2987–93. 10.1093/bioinformatics/btr509.

Li, Heng, and Richard Durbin. 2009. “Fast and Accurate Short Read Alignment with Burrows–Wheeler Transform.” Bioinformatics 25 (14): 1754–60. 10.1093/bioinformatics/btp324.

Liang, Zhikai, and James C. Schnable. 2016. “RNA-Seq Based Analysis of Population Structure within the Maize Inbred B73.” PLOS ONE 11 (6): e0157942. 10.1371/journal.pone.0157942.

Mamanova, Lira, Alison J Coffey, Carol E Scott, et al. 2010. “Target-Enrichment Strategies for next-Generation Sequencing.” Nature Methods 7 (2): 111–18. 10.1038/nmeth.1419.

McMullen, Michael D., Stephen Kresovich, Hector Sanchez Villeda, et al. 2009. “Genetic Properties of the Maize Nested Association Mapping Population.” Science (New York, N.Y.) (United States) 325 (5941): 737–40. 10.1126/science.1174320.

Michelmore, R. W., I. Paran, and R. V. Kesseli. 1991. “Identification of Markers Linked to Disease-Resistance Genes by Bulked Segregant Analysis: A Rapid Method to Detect Markers in Specific Genomic Regions by Using Segregating Populations.” Proceedings of the National Academy of Sciences of the United States of America (United States) 88 (21): 9828–32. 10.1073/pnas.88.21.9828.

Moore, Gordon E. 1965. “Cramming More Components onto Integrated Circuits.” Electronics 38 (8).

Multani, Dilbag S., Steven P. Briggs, Mark A. Chamberlin, Joshua J. Blakeslee, Angus S. Murphy, and Gurmukh S. Johal. 2003. “Loss of an MDR Transporter in Compact Stalks of Maize Br2 and Sorghum Dw3 Mutants.” Science (United States) 302 (5642): 81–84. 10.1126/science.1086072.

Neuffer, M. G., Guri Johal, M. T. Chang, and Sarah Hake. 2009. “Mutagenesis – the Key to Genetic Analysis.” In Handbook of Maize: Genetics and Genomics, edited by Jeffrey L. Bennetzen and Sarah Hake. Springer New York. 10.1007/978-0-387-77863-1_4.

Neuffer, M.G. 1988. “Designation of a Dominant Lax Midrib Mutant, Lxm1.” Maize Genetics Cooperation Newsletter 62: 53.

Neuffer, M.G. 1990. “Location and Description of Dominant Dwarf Mutants.” Maize Genetics Cooperation Newsletter 64: 51.

Neuffer, M.G. 1992. “Location and Designation of 8 Dominant Mutants from Chemical Mutagenesis and Spontaneous Origin.” Maize Genetics Cooperation Newsletter 66: 39–40.

Olukolu, Bode A., Guan-Feng Wang, Vijay Vontimitta, et al. 2014. “A Genome-Wide Association Study of the Maize Hypersensitive Defense Response Identifies Genes That Cluster in Related Pathways.” PLOS Genetics 10 (8): e1004562. 10.1371/journal.pgen.1004562.

Peiffer, Jason A., Maria C. Romay, Michael A. Gore, et al. 2014. “The Genetic Architecture of Maize Height.” Genetics (United States) 196 (4): 1337–56. 10.1534/genetics.113.159152.

Poland, Jesse A., Patrick J. Brown, Mark E. Sorrells, and Jean-Luc Jannink. 2012. “Development of High-Density Genetic Maps for Barley and Wheat Using a Novel Two-Enzyme Genotyping-by-Sequencing Approach.” PLOS ONE 7 (2): 1–8. 10.1371/journal.pone.0032253.

Quinlan, Aaron R., and Ira M. Hall. 2010. “BEDTools: A Flexible Suite of Utilities for Comparing Genomic Features.” Bioinformatics 26 (6): 841–42. 10.1093/bioinformatics/btq033.

Reid, L. M., K. Xiang, X. Zhu, B. R. Baum, and S. J. Molnar. 2011. “Genetic Diversity Analysis of 119 Canadian Maize Inbred Lines Based on Pedigree and Simple Sequence Repeat Markers.” Canadian Journal of Plant Science 91 (4): 651–61. 10.4141/cjps10198.

Saghai-Maroof, M. A., K. M. Soliman, R. A. Jorgensen, and R. W. Allard. 1984. “Ribosomal DNA Spacer-Length Polymorphisms in Barley: Mendelian Inheritance, Chromosomal Location, and Population Dynamics.” Proceedings of the National Academy of Sciences 81 (24): 8014–18. 10.1073/pnas.81.24.8014.

Sauer, Matt, Jianfei Zhao, Meeyeon Park, Rajdeep S Khangura, Brian P Dilkes, and R Scott Poethig. 2023. “Identification of the Teopod1, Teopod2, and Early Phase Change Genes in Maize.” G3 Genes|Genomes|Genetics 13 (10): jkad179. 10.1093/g3journal/jkad179.

Sawers, Ruairidh J. H., Joanne Viney, Phyllis R. Farmer, et al. 2006. “The Maize Oil Yellow1 (Oy1) Gene Encodes the I Subunit of Magnesium Chelatase.” Plant Molecular Biology (Netherlands) 60 (1): 95–106. 10.1007/s11103-005-2880-0.

Schichnes, Denise, Claudine Woo, and Michael Freeling. 1994. “The Lxm1 Gene Maps near Position 88 on 3L.” Maize Genetics Cooperation Newsletter 68: 16–17.

Sen, S., and G. A. Churchill. 2001. “A Statistical Framework for Quantitative Trait Mapping.” Genetics (United States) 159 (1): 371–87. 10.1093/genetics/159.1.371.

Shih, Shelly Y., Nikhil Bose, Anna Beatriz R. Gonçalves, Henry A. Erlich, and Cassandra D. Calloway. 2018. “Applications of Probe Capture Enrichment Next Generation Sequencing for Whole Mitochondrial Genome and 426 Nuclear SNPs for Forensically Challenging Samples.” Genes 9 (1). 10.3390/genes9010049.

Sindhu, Anoop, Satya Chintamanani, Amanda S. Brandt, Michael Zanis, Steven R. Scofield, and Gurmukh S. Johal. 2008. “A Guardian of Grasses: Specific Origin and Conservation of a Unique Disease-Resistance Gene in the Grass Lineage.” Proceedings of the National Academy of Sciences 105 (5): 1762–67. 10.1073/pnas.0711406105.

Singleton, W.R. 1959. “Height Potential in Brachytic-2 and Brachytic-3 Types.” Maize Genetics Cooperation Newsletter 33: 3–4.

Stolle, Eckart, and Robin F. A. Moritz. 2013. “RESTseq--Efficient Benchtop Population Genomics with RESTriction Fragment SEQuencing.” PloS One (United States) 8 (5): e63960. 10.1371/journal.pone.0063960.

Troyer, A. Forrest. 2004. “Background of U.S. Hybrid Corn II.” Crop Science 44 (2): 370–80. 10.2135/cropsci2004.3700.

Turner, Emily H., Choli Lee, Sarah B. Ng, Deborah A. Nickerson, and Jay Shendure. 2009. “Massively Parallel Exon Capture and Library-Free Resequencing across 16 Genomes.” Nature Methods (United States) 6 (5): 315–16. 10.1038/nmeth.f.248.

Wetterstrand, K. A. 2024. DNA Sequencing Costs: Data from the NHGRI Genome Sequencing Program (GSP). www.genome.gov/sequencingcostsdata.

Zhang, Xinyan, Meixia Zhao, Donald R McCarty, and Damon Lisch. 2020. “Transposable Elements Employ Distinct Integration Strategies with Respect to Transcriptional Landscapes in Eukaryotic Genomes.” Nucleic Acids Research 48 (12): 6685–98. 10.1093/nar/gkaa370.

